# bayNorm: Bayesian gene expression recovery, imputation and normalisation for single cell RNA-sequencing data

**DOI:** 10.1101/384586

**Authors:** Wenhao Tang, François Bertaux, Philipp Thomas, Claire Stefanelli, Malika Saint, Samuel Marguerat, Vahid Shahrezaei

## Abstract

Normalisation of single cell RNA sequencing (scRNA-seq) data is a prerequisite to their interpretation. The marked technical variability and high amounts of missing observations typical of scRNA-seq datasets make this task particularly challenging. Here, we introduce bayNorm, a novel Bayesian approach for scaling and inference of scRNA-seq counts. The method’s likelihood function follows a binomial model of mRNA capture, while priors are estimated from expression values across cells using an empirical Bayes approach. We demonstrate using publicly-available scRNA-seq datasets and simulated expression data that bayNorm allows robust imputation of missing values generating realistic transcript distributions that match single molecule FISH measurements. Moreover, by using priors informed by dataset structures, bayNorm improves accuracy and sensitivity of differential expression analysis and reduces batch effect compared to other existing methods. Altogether, bayNorm provides an efficient, integrated solution for global scaling normalisation, imputation and true count recovery of gene expression measurements from scRNA-seq data.

## Introduction

scRNA-seq is a method of choice for profiling gene expression heterogeneity genome-wide across tissues in health and disease^*1, 2*^. Because it relies on the detection of minute amounts of biological material, namely the RNA content of one single cell, scRNA-seq is characterised by unique and strong technical biases. These arise mainly because scRNA-seq library preparation protocols recover only a small fraction of the total RNA molecules present in each cell. As a result, scRNA-seq data are usually very sparse with many genes showing missing values (i.e. zero values, also called dropouts). The fraction of all transcripts recovered from a cell is called capture efficiency and varies from cell to cell, resulting in strong technical variability in transcripts expression levels and dropouts rates. Moreover, capture efficiencies tend to vary between experimental batches resulting in confounding “batch effects”. Correcting for these biases in order to recover scRNA-seq counts reflecting accurately the original numbers of transcripts present in a cell remains a major challenge in the field ^*3–5*^

A common approach to scRNA-seq normalisation is the use of cell-specific global scaling factors. These methods are based on principles developed for normalisation of bulk RNA-seq experiments and assume that gene specific biases are small^*3*^. Typically, read counts per cell are divided by a cell specific scaling factor estimated either from spike-in controls^*6*^, or directly from the transcriptome data using methods developed initially for bulk RNA-seq^*7–9*^ or specifically for scRNA-seq^*10, 11*^. A recent method called SCnorm extended the global scaling approach by introducing different scaling factors for different expression groups^*12*^.

Importantly, scaling methods do not correct for cell-to-cell variations in dropout rates, as genes with zero counts remain zero after division by a scaling factor. Several approaches have been designed to tackle this problem. A series of methods use zero-inflated distribution functions, to explicitly model the dropout charachteristics^*13–15*^. Alternatively, other studies have proposed to infer dropouts based on expression values pooled across cells or genes^*16–19*^. For instance, scImpute pools expression values across similar cell subpopulations in each dataset and imputes dropouts using a Gamma-Normal mixture model and population specific thresholds^*18*^. Similarly, the MAGIC package is based on pooling gene expression values across cells using a network-based similarity metric^*19*^. Another method is based on K-nearest neighbour smoothing, which uses Poisson distribution and aggregate information from similar cells^*20*^. Conversely, the SAVER approach pools expression values across genes within each cell using a Gamma-Poisson Bayesian model^*17*^. The Gamma-Poisson model is also used in two other packages called Splatter and scVI for simulating and normalising scRNA-seq data respectively^*21, 22*^. scVI belongs to new class of approaches which implement deep learning variational autoencoder or autoencoder methods^*16, 21, 23–25*^. For instance, DCA, an autoencoder method, utilises a zero-inflated negative binomial noise model^*16*^. Experimental batch-to-batch variations are another common source of technical variability in scRNA-seq data. The origin of batch effects is not fully understood but results at least in part from differences in average capture efficiencies across experiments^*26*^. Several methods have been developed to specifically remove batch effect in scRNA-seq data^*27–29*^.

The methods discussed above, treat normalisation, imputation, and batch effect correction as separate tasks. Moreover, they rely on strong assumptions such as the zero-inflation model. Here we provide a detailed account of a novel integrated approach called bayNorm which performs all the processing steps discussed above at the same time using minimal assumptions. We compared its performance with a series of available packages focusing on true count recovery, differential expression analysis and batch effect correction.

### The bayNorm rationale

bayNorm is a Bayesian implementation of global scaling normalisation that simultaneously imputes missing values in scRNA-seq data. bayNorm generates for each gene (*i*) in each cell (*j*) a posterior distribution of original expression counts 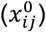, given the observed scRNA-seq read count for that gene (*x*_*ij*_) (**Fig. 1a**). Using the Bayes rule we have:

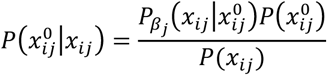

**Figure 1:**
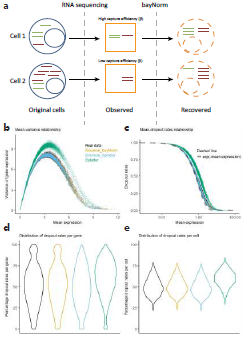
A binomial model of mRNA capture is consistent with the statistics of raw experimental scRNA-seq data. (a) Cartoon illustration of the bayNorm approach. Only a fraction of the total number of mRNAs present in the cell is captured during scRNA-seq library preparation. This occurs with a global probability called capture efficiency (*β*). Using cell-specific estimates of *β*, bayNorm aims at recovering the original number of mRNA of each gene present in each cell. (b)-(e) Comparisons between raw experimental scRNA-seq data from the Klein study[1] and synthetic data obtained using the Binomial bayNorm (orange), Binomial Splatter (blue), or Splatter[2] (green) simulation protocols (Supplementary Note 1). (b) Variance vs mean expression relationship. (c) Dropout rates vs mean expression relationship (note that Binomial Splatter and Binomial bayNorm are on top of each other in this panel). The dotted line shows the *e*^(-Mean expression)^ function. (d) Distribution of dropout values per gene. (e) Distribution of dropout values per cell.

Where 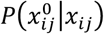 is the posterior distribution of true gene expression counts of a given gene in a given cell. 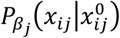 is a likelihood function that depends on the cell specific capture efficiency (*β*_*j*_). Specific capture efficiencies can be estimated using spike-in controls or directly from the data using scaling factors provided by different methods^*3*^ and normalised to the dataset’s mean capture efficiency <*β*> (see Methods).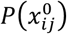 is a gene specific prior expression distribution and (*x*_*ij*_) is the marginal likelihood. The outputs of bayNorm are either samples (3D array) or point estimates (2D array) from the posterior distributions (**Fig. S1**).

### The binomial model is an appropriate choice for the bayNorm likelihood function

The bayNorm likelihood function 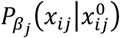 is at the core of the approach and describes the empirical distribution of the raw experimental scRNA-seq counts. The binomial model describes the random sampling of a fraction of a cell transcriptome with constant probability. This is a simple model of transcript capture in scRNA-seq^*30*^ and we therefore hypothesised that it would be a good choice for bayNorm likelihood function. For the prior 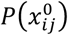 we assume a negative binomial model, which describes the bursty distribution of mRNAs in simple models of gene expression ^*31, 32*^. Gene specific prior parameters are estimated using an empirical Bayes approach by pooling gene expression values across multiple cells of the dataset (see Methods for details).

To validate our choice of binomial likelihood model and prior estimates, we generated simulated scRNA-seq data based on these assumptions and investigated how closely they captured statistics of several published scRNA-seq datasets (**Fig. 1 b-e, Fig. S2-7**)^*12, 30, 33, 34*^. The simulations assumed mRNA counts per cell that followed negative binomial distributions and used gene specific priors obtained with bayNorm (**Fig. 1**, Binomial_bayNorm), or sampled from estimates obtained with a modified version of the Splatter package (**Fig. 1**, ‘Binomial_Splatter’, Supplementary Notes 1)^*22*^. These were compared with simulations generated with the original Splatter package which is based on the Gamma-Poisson distribution^*22*^. We note that in Splatter, scaling factors are multiplicative to the Gamma distribution’s mean. In bayNorm, however, the cell specific capture efficiencies, which act as scaling factors, are set as the probability parameter of the binomial model. We found that the binomial model captures the variance-mean relationship of experimental scRNA-seq data well (**Fig. 1b**).

Another important feature of scRNA-seq data is their large amount of missing values, or dropouts, and several models have been proposed to explain this phenomenon^*14, 15, 26, 35, 36*^. We therefore investigated how well the binomial model would capture dropout rates in experimental data. Our simulated dataset generated using the ‘Binomial_bayNorm’function reproduced accurately the dependence of dropout fractions on gene expression means performing better than Splatter (**Fig. 1c-e**). Moreover, a parameter free approximation based on the binomial model predicted the dropout fraction to depend on an exponential of the negative mean expression (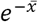, see Methods). This functions produced a very close fit to the experimental data providing additional support for our choice of the binomial model (**Fig. 1c**). Notably, the Binomial_bayNorm simulation protocol using inferred gene-specific priors together with cell specific parameters *(β*_*j*_) was the only one that recovered the distribution of dropout rates per gene observed in experimental data (**Fig. 1d**). Finally, the results presented on **Fig. 1b-e** could be replicated consistently using several additional experimental scRNA-seq datasets (**Fig. S2-7**).

The datasets discussed so far include sequencing scores corrected for PCR amplification biases using unique molecular identifiers (UMIs)^*37*^. Some popular protocols, however, do not include UMIs, and are therefore likely to be less well described by the binomial distribution due to technical variability arising from PCR amplification bias. Accordingly, their dependence of dropout fractions on the mean expression has been reported to be more complex than in UMI-based datasets ^*36*^. We investigated this issue further and found that a simple scaling of non-UMI raw data by a constant factor produced a reasonable match to the binomial model (**Fig. S9**; see Methods). This scaling factor can be interpreted as the average number of times original mRNA molecules were sequenced after PCR amplification. This indicates that, provided appropriate scaling, non-UMI datasets are also compatible with the bayNorm model. Importantly, as bayNorm recovered dropouts rates successfully in both UMI-based and non-UMI protocols without the need of specific assumptions, we conclude that invoking zero-inflation models is not required to describe scRNA-seq data. Consistent with this, the differences in mean expression levels of lowly expressed genes observed between bulk and scRNA-seq data, which were suggested to be indicative of zero-inflation, were recovered by our simulated data using the binomial model only (**Fig. S10**)^*38*^.

We note that the ability of simulation protocols to recover the statistics of experimental data depended intimately on the value of cell-specific capture efficiencies (*β*_*j*_). We used different ways to estimate *β* (spike-in, Scran scaling factors, trimmed means, or housekeeping genes; Supplementary Note) together with different <*β>*in the Binomial_Splatter simulation protocol. We found that changes in *β* values affected recovery of the distribution of dropout rates per cell. (**Fig. S8**). In particular, we found that the use of spike-in controls or of housekeeping reference gene expression levels did not improve estimates of capture efficiencies (**Fig. S8c-f**). Altogether, this analysis demonstrates that accurate statistics of experimental scRNA-seq data can be consistently retrieved using the binomial model and empirical Bayes estimation of gene expression parameters implemented in bayNorm along with accurate estimates of cell-specific capture efficiencies.

### bayNorm enables recovery of true gene expression distributions from scRNA-seq data

Single-cell RNA-seq provides a unique opportunity to study stochastic cell-to-cell variability in gene expression at a near genome-wide scale. However, doing this requires normalisation approaches able to retrieve from scRNA-seq data transcripts levels matching quantitatively *in vivo* mRNA numbers^*33*^. With this in mind, we evaluated bayNorm performance in reconstructing true gene expression levels from a series of experimental scRNA-seq datasets that contained matched single molecule fluorescence *in situ* hybridisation (smFISH) measurements for a series of genes. We used global mean capture efficiencies <*β>* estimated directly from smFISH together with gene specific priors informed by the sequencing data (**Fig S11**). After bayNorm normalisation, scRNA-seq counts reproduced accurately count distributions obtained by smFISH for several mRNAs (**Fig 2a-b**). We then compared bayNorm performance with a series of published normalisation methods (Supplementary note 4, **Fig 2**). All methods captured mean smFISH counts across different genes well (**Fig. 2c-d, Fig S11**). However, noise in gene expression (coefficient of variation, CV) and expression dispersion (Gini coefficient) measured by smFISH were better captured by bayNorm compared to normalisation by scaling or by several recent normalisation and imputation methods (**Fig. 2e-f, Fig. 2g-h**) ^*12, 16–19*^. bayNorm’s good performance could also be confirmed in a series of simulation studies (**Fig S12**, Supplementary note 1). In summary, bayNorm combined with gene specific priors inferred directly from the scRNA-seq data, retrieves gene expression variability matching closely smFISH data.

**Figure 2:**
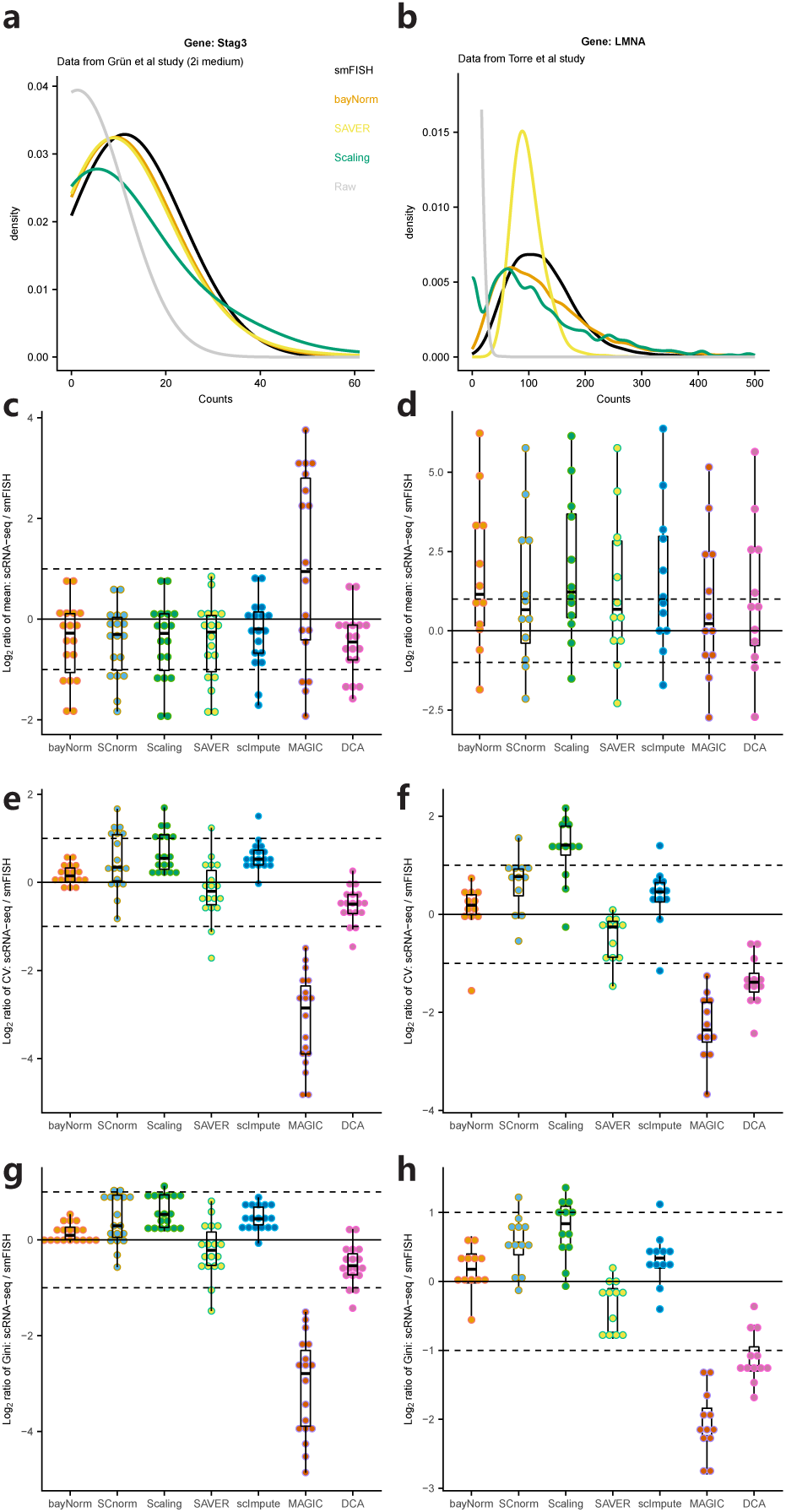
bayNorm recovers distributions of gene expression observed by smFISH. (a) Stag3 mRNA distribution for cells grown in 2i measured by smFISH or by scRNA-seq and normalised with different methods (from Grn study). Raw denotes unnormalised scRNA-seq data. (b) As in (a) for the LMNA gene (from Torre study). Legend as in (a). (c) Log_2_ ratio between the means of scRNA-seq measurements for 18 genes normalised by different methods and their matched smFISH measurements (from Grn study). (d) As in (c) using 12 genes (Torre study). (e) Log_2_ ratio between the CV of scRNA-seq measurements for 18 genes normalised by different methods and their matched smFISH measurements (from Grn study). (f) As in (e) using 12 genes (from Torre study). (g) Log_2_ ratio between the Gini coefficients of scRNA-seq measurements for 18 genes normalised by different methods and their matched smFISH measurements (from Grn study). (h) As in (c) using 12 genes (from Torre study). For the bayNorm and SAVER normalised datasets, 20 or 5 samples were generated from posterior distributions for the Grn and the Torre studies, respectively. For bayNorm and SAVER, normalized counts across cells and samples are used. All normalised datasets except bayNorm and the Scaling method have been divided by the *< β >* value used in bayNorm procedure. For this analysis smFish data were normalised for variation in total transcript numbers using either cell size measurements (Grn study) or expression levels of a house keeping gene (Torre study) as detailed in Supplementary note 3.

### bayNorm enables accurate and sensitive differential expression analysis

Differential genes expression analysis (DE) in scRNA-seq studies is challenging as several factors including variability in capture efficiencies, dropout rates, sequencing depth, and experimental batch effects can introduce significant, yet spurious, differential expression signal. Normalisation and imputation approaches have, therefore, a significant impact on the sensitivity and accuracy of DE analysis protocols. Two features of the bayNorm approach have the potential to improve the performance of DE analysis. Firstly, bayNorm posterior distribution of original counts maintains the uncertainty resulting from small capture efficiencies and could therefore reduce false positive DE discovery rates^*39*^. Secondly, the use of priors specific to each group of cells compared in the DE analysis could increase true positive discovery rates. With this in mind, we have assessed bayNorm performance in DE analysis using several experimental scRNA-seq datasets and compared it to other existing methods. To identify DE genes we use MAST^*13*^, which performs well in terms of false positives rates, precision and recall^*40*^. MAST was first applied to individual sample from the bayNorm posterior distribution (3D array, **Fig. S1**). Differentially expressed genes were then called based on the median of Benjamini-Hochberg adjusted P-values of the individual samples^*28*^.

As mentioned above, differences in capture efficiencies between cells is a source of technical variability that could affect DE analysis. To test bayNorm’s ability to correct for this bias, we selected the 100 cells with the highest and lowest capture efficiencies based on total counts in a recent UMI-based scRNA-seq study^*30*^. We then applied bayNorm to the 200 cells using global prior estimation based on the combination of the two groups (see Methods). In this design, the two groups of cells differ based only on their capture efficiencies, and significant differential expression is therefore not expected. **Fig. 3a** shows the number of genes called differentially expressed as a function of increasing average expression levels using a series of normalisation and imputation methods^*12*^. bayNorm normalised data show almost no differentially expressed genes, outperforming all the other methods. Moreover, log_2_ gene expression ratios between cells of the two groups, were consistently close to zero, confirming bayNorm ability to correct for biases inherent to different capture efficiencies in UMI-based datasets (**Fig. 3b**).

**Figure 3:**
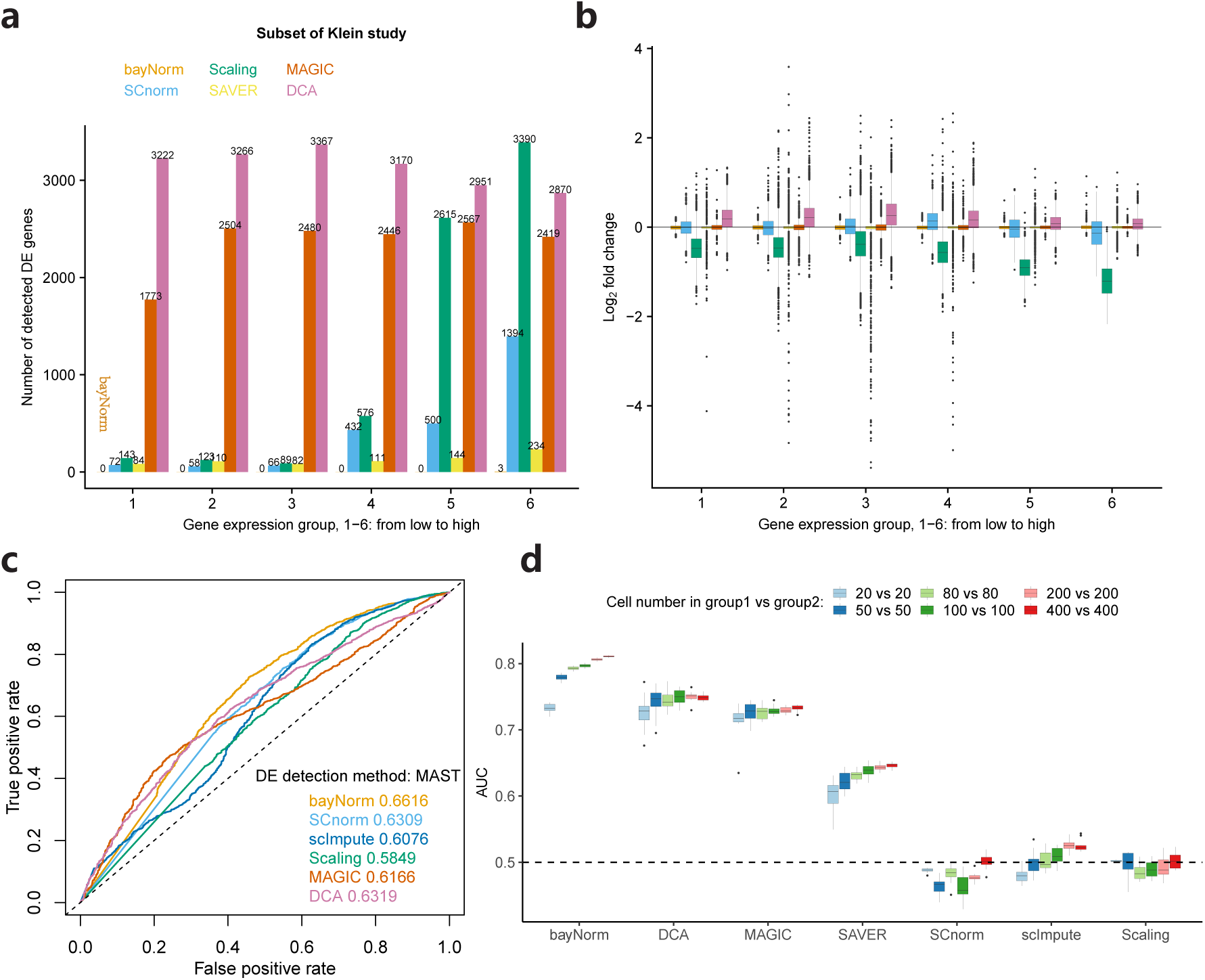
bayNorm enables robust and sensitive differential expression analysis. (a) Number of differentially expressed genes between the 100 cells with the highest and the 100 cells with the lowest total counts in (Klein study). DE genes were called using the MAST package (*P*_MAST_ *<* 0.05) and plotted for 6 groups of genes with increasing mean expression (1-low 6-high). (b) Log_2_ fold change from (a). (c) Differential expression analysis using MAST for different normalization methods (Islam study) using a benchmark list of DE genes obtained from matched bulk RNA-seq data[3]. (d) Differential expression analysis using data from Soumillon study[4]. Ten samples of 20, 50, 80, 100, 200 or 400 cells were selected randomly from two groups of stage-3 differentiated cells at day 0 (D3T0) or day 7 (D3T7). DE detection was performed between groups as described at the top of the figure using a list of DE genes obtained from matched bulk RNA-seq data as a benchmark (1000 genes with the largest magnitude of log fold-change between the D3T0 and D3T7 samples)[3].

Sequencing depth is another parameter affecting DE analysis especially because it impacts on the dropout rates of lowly expressed genes. Moreover, differences in sequencing depth are likely to affect levels of capture efficiencies, especially for non-UMI datasets where PCR biases are not accounted for. To assess bayNorm’s ability to correct for this source of bias, we used a benchmark dataset published by Bacher and colleagues^*12*^ that consists of non-UMI based scRNA-seq data for two groups of cells isolated from a single culture and sequenced to a depth of either 1 million or 4 million reads per cell. bayNorm and other imputation methods performed well in this setting (**Fig. S13**). However, a global scaling approach on its own led to poor results, unless performed independently on groups of genes with similar mRNA expression levels as in SCnorm. Finally, bayNorm corrected robustly for variability in sequencing depth when applied to a series of simulated datasets (**Fig. S14-15**)^*12*^.

We have shown that bayNorm is efficient at removing spurious differential expression from scRNA-seq data caused by variability in capture efficiencies and sequencing depth. We next explored bayNorm performance in supporting sensitive and robust detection of genes truly regulated between samples. To do this, we used two experimental scRNA-seq datasets^*41, 42*^ and lists of benchmark DE genes derived from matched bulk RNA-seq data^*40, 43*^. To maximise sensitivity, we used priors specific to each groups of cells in the comparison (we call this design “local priors”). With the first dataset, bayNorm normalised data generated an AUC value as high as other normalisation methods demonstrating that the approach supports sensitive DE detection (**Fig. 3c**). Analysis of the second dataset (UMI-based)^*42*^ confirmed this observation with bayNorm performing better than all other methods (**Fig. 3d**). Importantly, bayNorm performance did not depend on the number of cells in each group, except for groups with very low numbers of cells (**Fig. 3d**, **Fig. S16**). Finally, using a series of simulated datasets, we explored situations where the compared groups have different mean capture efficiencies and found that bayNorm supported robust DE detection in all cases (**Fig. S17**).

Three important parameters should be considered before bayNorm normalisation: i) the choice of priors, ii) the choice of average capture efficiencies <*β>*, iii) the choice of bayNorm output format. Prior parameters can be either estimated for all cells across groups (global) or within each group (local). Since priors are gene specific, applying bayNorm across homogeneous cells (*i.e.* using global prior) allows for mitigating technical variations (**Fig S18a-b**). On the other hand, using priors estimated “locally” within each group amplifies differences in signals between heterogeneous groups of cells increasing sensitivity (**Fig S18c-d**). Average capture efficiencies <*β>* are specific to each scRNA-seq protocol and reflect their overall sensitivity. This value represents the ratio of the average number of mRNA molecules sequenced per cell to the total number of mRNA molecules present in an average cell. It is not always easy to determine as quantitative calibration methods such as smFISH are not widely used, and approaches based on spike-in controls have important shortcomings^*3*^. We investigated the impact of inaccurate estimation of <*β>* on biases in DE detection. Critically we found that DE results based on bayNorm normalised data are not affected significantly by a 2 fold change of <*β>* (**Fig. S20-S21**). Finally, bayNorm output consists of either samples from its posterior distributions (3D array) or the modes of these distributions as point estimates (2D arrays). For DE analysis using MAST, 3D arrays reduces false positive rates but 2D arrays perform slightly better in terms of AUC (**Fig S18c-d**). **Fig S19** shows DE results for two other non-parametric methods: ROTS^*44*^ and Wilcoxon test^*40*^. Both approaches perform equally well with 3D arrays but show variable results when applied to 2D arrays with the Wilcoxon test performing less well.

In summary, our analysis demonstrates that in addition to correcting for technical biases, bayNorm also supports robust and accurate DE analysis of a wide range of experimental and simulated scRNA-seq datasets.

### bayNorm correction of experimental batch effects

scRNA-seq protocols are subject to significant experimental batch effects^*34*^. In cases where the study design does not take this problem into account by distributing cases and controls across batches for instance, batch effects can lead to artefactual differences in gene expression of single cells, resulting in inaccurate biological conclusions. bayNorm can mitigate batch effects in two ways. First, as described above, bayNorm efficiently corrects for differences in capture efficiencies which is a pervasive source of batch-to-batch variability^*38*^. Second, the use of bayNorm data-informed priors is an efficient way to mitigate batch variation by estimating prior parameters across different batches but within the same biological condition. To investigate bayNorm’s performance for batch effect correction we use data from the Tung study ^*34*^ where scRNA-seq data were obtained in triplicates for three induced pluripotent stem cell lines (iPSC) derived from three individuals. Sequencing libraries were prepared in three experimental batches, each containing one repeat of each line^*34*^. We first used priors calculated within each individual, but across batches (bayNorm local (individual)). This strategy allows for maintaining differences between individuals while minimising batch effects as illustrated by PCA analysis (**Fig. 4a-b, Fig S22**). To assess the normalisation performance quantitatively, we extracted the number of genes differentially expressed between each pair of batches within the same individual (**Fig S23**). We defined the ratio of the number of DE genes (adjusted P_MAST_ < 0.05) and the total number of genes (13058) to be the false positive rates (FPR). In theory, batch effects should be the main source of differential expression between these samples^*34*^. In parallel, we tested whether bayNorm also maintained differences between individuals using the same settings. To do this, we defined DE genes between the iPSC lines NA19101 and NA19239 and compared it to a benchmark list of 498 DE genes^*43*^. Efficient batch effect correction is expected to minimise FPR while maximizing Area Under the Curve (AUC) values of DE detection between individuals. We find that using bayNorm with “within individual” local priors (estimated across different batches within the same line) outperformed other methods in terms of correcting batch effects while maintaining meaningful biological information. As expected, using bayNorm and global priors (estimated across batches and individuals, bayNorm global) preserves low FPR, but reduces AUC significantly. Finally, using bayNorm with “within batch” local priors (bayNorm local (batch)) result in higher false positive rates, which is also expected.

**Figure 4:**
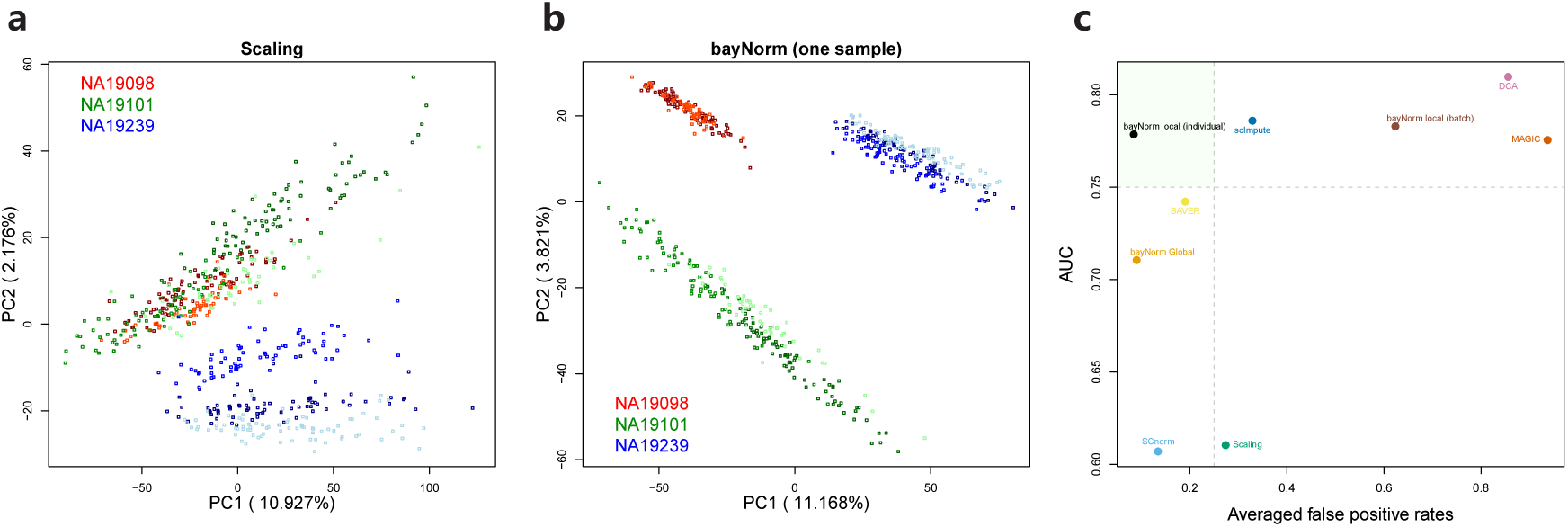
bayNorm normalisation reduces experimental batch effects. (a) PCA plots of data from the Tung study normalised using global scaling. Each colour represent a different cell line derived from a different individual. Colour shades represent different batches within a line/individual. (b) As in (a) using bayNorm normalization. (c) Differentially expressed genes were called between lines NA19101 and NA19239. DE genes from matched bulk RNA sequencing data were used as a benchmark set and AUC values were calculated. In parallel, DE genes were called between different batches within each line (7 pair of comparisons in total). The ratio of the number of DE genes per line and the total number of genes (13058) between batched is defined as the DE false positive rate (FDR). FDRs were averaged across the 7 pairs. Each normalisation method results in a pair of averaged FDR and AUC values that is displayed on the figure. Normalisation methods are colour-coded. The vertical and horizontal dashed lines represent 0.25 and 0.75 indicative cutoffs respectively. bayNorm was applied either across batches but within lines (“bayNorm local (individual)”) or across all cells (“bayNorm global”) or within each batch (“bayNorm local (batch)”).

Overall we have showed that the flexibility of priors selection afforded by bayNorm Bayesian approach enables robust correction of batch effects, while maintaining sensitive detection of differentially expressed genes.

### Conclusions

We introduced bayNorm, a versatile Bayesian approach for implementing global scaling that simultaneously provides imputation of missing values and true counts recovery of scRNA-seq data. Bayesian methods have been applied to different aspects of RNA-seq data analysis before^*11, 15, 17, 45*^. The approach most related to bayNorm is taken by SAVER which uses a Poisson-Gamma model and pooling information across genes for true-count recovery^*17*^. In contrast, bayNorm uses a binomial model of mRNA capture as likelihood and achieves similar or improved performance relative to SAVER on real and simulated data (Fig 2-4). We showed that using the binomial model and an empirical Bayes approach to estimating gene expression priors across cells results in simulated data almost identical to experimental scRNA-seq measurements. Importantly, this suggests that zero-inflated models are not required to explain the frequency of dropout observed in scRNA-seq. Although designed initially for UMI-containing scRNA-seq protocols, a simple scaling factor makes bayNorm applicable to non-UMI data as well. This flexibility will allow using this approach with most present and future scRNA-seq datasets. We showed using datasets that combine smFISH and scRNA-seq, that bayNorm is accurately recovering true gene expression across a wide range of expression levels. This approach could therefore be particularly useful for quantitative analysis of more difficult scRNA-seq datasets, such as those generated from small quiescent cells or microbes, for instance. In fact, we have recently used bayNorm successfully in the first scRNA-seq study of fission yeast^*46*^. One of the most powerful features of bayNorm is its use of gene expression priors directly calculated from gene expression values across cells. We showed that by grouping cells according to experiment design or phenotypic features increased significantly the robustness and sensitivity of differential expression analysis. This allows almost complete removal of sequencing depth and capture efficiency biases, and reduced batch effects. Critically, this approach preserved accurate and sensitive detection of benchmark DE genes.

Accurate estimation of cell capture efficiencies (or scaling factors) is central to most scRNA-seq normalisation methods including bayNorm. Interestingly, we observed that the choice of cell specific capture efficiencies affect how closely simulated data recovers statistics of real data. We therefore propose that comparison of drop-out rates per cell in simulated datasets and experimental data could be used as a tool to inform appropriate choice of global scaling factors and mean capture efficiency estimates. The option to tailor bayNorm priors based on phenotypic information about cell subpopulations will be a powerful asset for discovery of gene expression programmes associated with specific phenotypic features of single cells such as cell size^*46*^. Finally, the concepts and mathematical framework behind bayNorm will be useful if combined with other emerging theoretical approaches such as deep learning, for instance ^*16, 21, 23–25*^. Overall, bayNorm provides a simple and integrated solution to remove the technical biases typical of scRNA-seq approaches, while enabling robust and accurate detection of cell-specific changes in gene expression. bayNorm has been made freely available as an R package (see Methods), and will be submitted to Bioconductor.

## Acknowledgements

We are grateful to Dan Hebenstreit for critical reading of the manuscript. The benchmark lists used in the Islam and Tung studies were kindly provided by Maria K. Jaakkola and Chengzhong Ye respectively. We would like to thank Rhonda Bacher for providing R code for running MAST and producing figures 3a and 3b. We would like to thank Mo Huang for the code for preprocessing data from the Torre Study. We would also like to thank Lennart Kester for providing smFISH data used in the Grün study. This research was supported by the UK Medical Research Council, and a Leverhulme Research Project Grant (RPG-2014-408). WT is supported by a Roth Scholarship from the Department of Mathematics at Imperial College. PT acknowledges a fellowship from The Royal Commission for the Exhibition of 1851. The authors used the computing resources of the UK Medical Bioinformatics partnership (UK MED-BIO; aggregation, integration, visualisation and analysis of large, complex data), which is supported by the UK Medical Research Council (grant no. MR/L01632X/1) and the Imperial College Research Computing Service (DOI: 10.14469/hpc/2232) for access to their HPC facilities (CX1 cluster).

## Methods

### 1 The Bayesian model used in bayNorm

A scRNAseq dataset is typically represented in a matrix of dimension *P × Q*, where P denotes the total number of genes observed and Q denotes the total number of cells studied. The element *x*_*ij*_ (*i* ∈{1, 2, *…, P}* and *j ∈ {*1, 2, *…, Q}*) in the matrix represents the number of transcripts reported for the *i*^th^ gene in the *j*^th^ cell. This is equal to the total number of sequencing reads mapping to that gene in that cell for a non-UMI protocol. For UMI based protocols this is equal to the number of individual UMIs mapping to each gene [5, 6]. The matrix can include data from different groups or batches of cells, representing different biological conditions. This can be represented as a vector of labels for the cell groups or conditions (*C*_*j*_).

A common approach for normalizing scRNAseq data is based on the use of a global scaling factor (*s*_*j*_), ignoring any gene specific biases (for a recent review see[7]). The normalized data 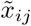 is obtained by dividing the raw data for each cell *j* by the its global scaling factor *s*_*j*_:

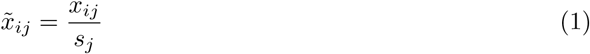

In bayNorm, we implement global scaling using a Bayesian approach. We assume given the original number of transcripts in the cell 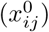, the number of transcripts observed (*x*_*ij*_) follows a Binomial model with probability *β*_*j*_ [1], which we refer to as capture effeiciency and it represents the probability of original transcripts in the cell to be observed. In addition, we assume that the original number or true count of the *i*^th^ gene in the *j*^th^ cell 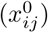 follows Negative Binomial distribution with parameters mean (*µ*), size (or dispersion parameter, *φ*), such that:

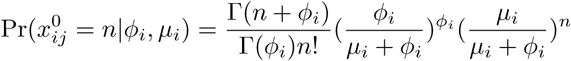

So, overall we have the following model:

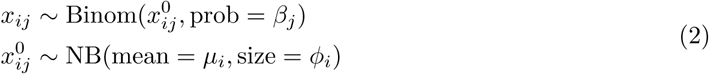

Using the Bayes rule, we have the following posterior distribution of original number of mRNAs for each gene in each cell:

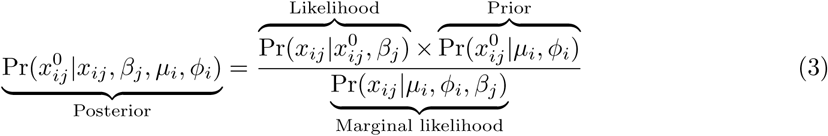

The prior parameters *µ* and *φ* of each gene were estimated using an empirical Bayesian method as discussed in detail in Section 4 below.

The marginal distribution for gene *i* in cell *j* is

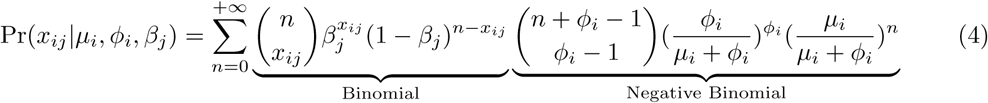

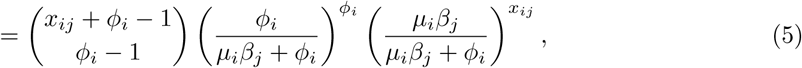

which follows from using

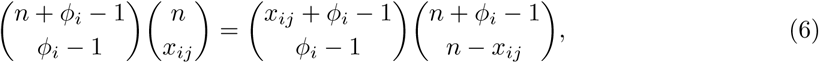

and

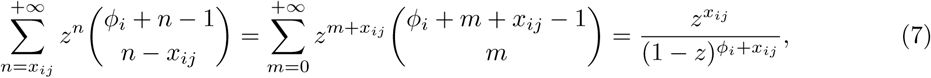

with 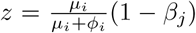in Eq. (4). Hence we have that the number of transcripts reported for the *i*^th^ gene in the *j*^th^ cell

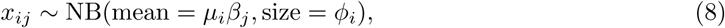

has a Negative Binomial distribution with mean *µ*_*i*_*β*_*j*_ and size *φ*_*i*_.

It can also be shown that the posterior distribution of 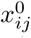 is a shifted Negative Binomial distribution.To sample from the posterior distribution, we note that the original count can be expressed as

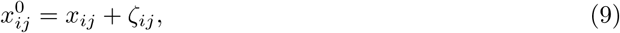

where *ζ*_*ij*_ is the *lost* count satisfying

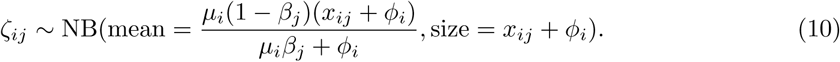

The posterior mean and variance then evaluate to

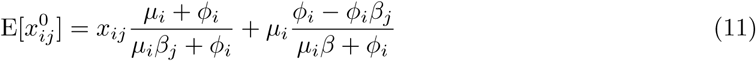

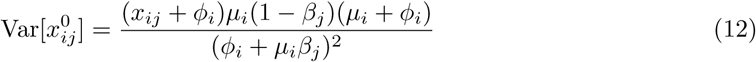

Note that when *φ*_*i*_ is small, the mean of posterior tends to 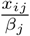. After estimating the posterior distribution for each gene in each cell, we can either sample a certain number of draws from it (3D array output, see Supplementary Figure S1) or extract the mean or mode of posterior [8] as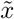 (2D array output, see Supplementary Figure S1).

### 2 Binomial distribution and dropout probability

The binomial model of capture in scRNA-seq predicts the dropout rate for a particular gene:

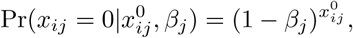

in a given cell *j*. Across a group of non-homogeneous cells, we may approximate this expression b0079

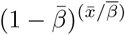

For small 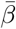 this expression tends to 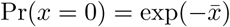 In dropout vs mean expression (dropout-mean) (Figure 1c, Sup Figures S2c, S3c, S4c, S5c, S6c and S7c), the line 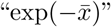 follows the lower limit of the trend. We note that a Poisson model of RNA-seq that is used by several authors also predicts dropout rates to be Pr(*x* = 0) = *λ*^0^*/*0! exp(*-λ*) = exp(*-λ*), where 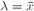 [9, 2, 10].

To further show that Binomial distribution can capture the relationship between dropout rates and mean expression, we simulated data based on real experimental data[1, 11, 12] by adapting simulation protocols proposed in the R package Splatter[2]. The details about the simulation procedure can be found in the supplementary note 1. The resulting dropout-mean plot of simulated data based on Binomial model is very close to that of the real scRNA-seq data for UMI-based protocols. As shown in the Supplementary Figures S2c, S3c, S4c, S5c and S6c, the dropout-mean trend of UMI data is close to the asymptotic line 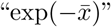 (Binomial Splatter and Binomial bayNorm simulated data perform similar to each other and the real experimental data). data based on real experimental data[1, 11, 12] as discussed in the results and supplementary note 1. The resulting dropout-mean plot of simulated data based on Binomial model is very close to that of the real scRNA-seq data for UMI-based protocols.

### 3 Estimation of capture efficiencies

Cell specific capture efficiency *β*_*j*_ and global scaling factor (*s*_*j*_) are closely related. We can transform scaling factors estimated by different methods (see below) into *β*_*j*_ values with the following formula:

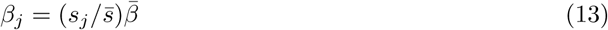

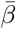 a scalar, is an estimate of global mean capture efficiency across all cells, which ranges between 0 and 1.

There are two different methods for estimating 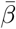 and *β*_*j*_:

1. If spikeins or smFISH data are available they can be used to estimate capture efficiencies. We can either divide the total number of observed spikins in each cell by the total number of input spike-ins, or we can fit a linear regression[1] to estimate the cell specific *β*_*j*_. If smFISH data is available, we can fit a linear regression between the mean expression of raw data (response variable) and the mean expression of the smFISH data (explanatory variable). The coefficient of the explanatory variable can be used as *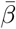 [13]*
2. The raw data itself can be directly used for estimation of cell specific global scaling factors (s_j_). Then equation 13 and an estimate of 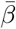 can be used to estimate *β*_*j*_. There are different methods available for estimation of global scaling factors. Some were developed for bulk RNA-seq data[14, 15] and some are specific to scRNA-seq data[16, 17]. The value of 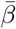 depends on the protocol used and can be batch dependent. For example, for Droplet based protocol, it is about 0.06[1] or 0.12[18]. 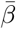 can also be estimated by spike-ins or smFISH data as explained above.

We finally note, that estimates of capture efficiency discussed above will assume cells have simular original transcript content. Therefore, the bayNorm outputs estimates of original transcript counts for a typical cell, which is corrected for variation in cell size and transcript content. This is usually desirable for down-stream analysis such as DE detection. However, if one is interested in absolute origianl count and has additional information such as cell size or total transcirpt content per cell, the capture efficiencies can be approporiatly rescaled for this purpose.

### 4 Estimation of prior parameters

#### 4.1 Maximisation of marginal distribution

Using an emperical bayes approach, one can use the maximisation of marginal likelihood distribution of the observed counts across cells to estimate prior parameters [19]. Let *M*_*i*_ denotes the marginal likelihood function for the *i*^th^ gene across cells. Assuming independence between cells, the log-marginal distribution for the *i*^th^ gene is

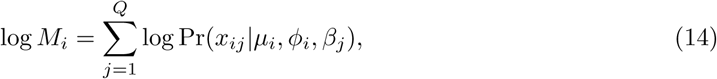

WherePr(*x*_*ij*_|*|µ*_*i*_,*φ*_*i*_, *β*_*j*_) is the Negative Binomial in Eq. (5). Maximizing of Eq. (14) yields the pair (*µ*_*i*_, *φ*_*i*_).

The above optimization needs to be done for each of the P genes. We refer to the *φ* and/or *µ* estimated by maximizing marginal distribution as BB estimates for convenience, because bayNorm utilizes spectral projected gradient method (spg) from the R package named “BB”. Optimizing the marginal distribution with respect to both *µ* and *φ* (2D optimization) is computationally intensive. If we had a good estimate *µ*, then we could optimize the marginal distribution with respect to *φ* alone, which would be much more efficient.

#### 4.2 Method of Moments

A heuristic way of estimating *µ*_*i*_ and *φ*_*i*_ is through a variant of the Method of Moments. The first step is to do a simple normalization of the raw data, to scale expressions given the cell specific capture efficiencies (*β*_*j*_). The simple normalized count 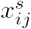 is calculated as following:

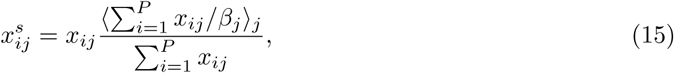

where the numerator of the scaling factor of *x*_*ij*_ is obtained by taking the average of scaled total counts across cells.

Based on simple normalized data, we are able to estimate prior parameters *µ* and *φ* of the Negative Binomial distribution using the Method of Moments Estimation (MME), which simply equates the theoretical and empirical moments. This estimation method is fast and simulations suggests it provides good estimates of *µ* but the drawback is that the estimation of *φ* show a systematic bias (see Supplementary Figure S24 a-b).

#### 4.3 The combined method

Based on simulation studies (Supplementary Figure S24), the most robust and efficient estimation of *µ* and *φ* can be obtained using the following combined approach, which is the default setting in bayNorm:

1. Based on simple normalized data, we use the MME method for each gene to obtain MME estimated *µ* and *φ*.
2. Although the BB estimated *φ* is much closer to the true *φ*, many estimates are at the upper boundary of the search space (Supplementary Figures S24 c-d). So, we find adjusting the MME estimated *φ* by a factor which can be estimated by fitting a linear regression between MME estimated *φ* and BB estimated *φ* works best (Supplementary Figures S24 c-d). This adjusted MME estimated *φ* together with the MME estimated *µ* and estimates of *β*_*j*_ can be used in approximating posterior distribution for each gene in each cell.

Cells are grouped together for prior estimation, based on cell-specific attributes (*C*_*j*_). Prior estimation can be done over all cells irrespective of the experimental condition. We refer to this procedure as “global”. Alternatively, suppose that there are multiple groups of cells in the datasets and we have reasons to believe each group could behave differently. Then we can estimate the prior parameters “*µ* and *φ*” within each group respectively (within groups with the same *C*_*j*_ value). We refer to this procedure as “local”. Estimating prior parameters across a certain group of cells based on “global” procedure allow for removing potential batch effects. Multiple groups normalization based on “local” procedure allows for amplifying the inter-groups’ differences while mitigating the intra-group’s variability, which is suitable for DE detection.

### 5 Code availability

The R package bayNorm is available at https://github.com/WT215/bayNorm.

The codes for producing figures in the paper are provided at https://github.com/WT215/bayNorm_papercode.

In the Bacher study, the code for running MAST and log fold change calculation was kindly provided by Rhonda Bacher, the author of SCnorm[20].

In the Torre study, the code for transforming counts per million normalized data to UMI data was kindly provided by Mo Huang, the author of SAVER[9].

## Supplementary figures

**Supplementary Figure 1:**
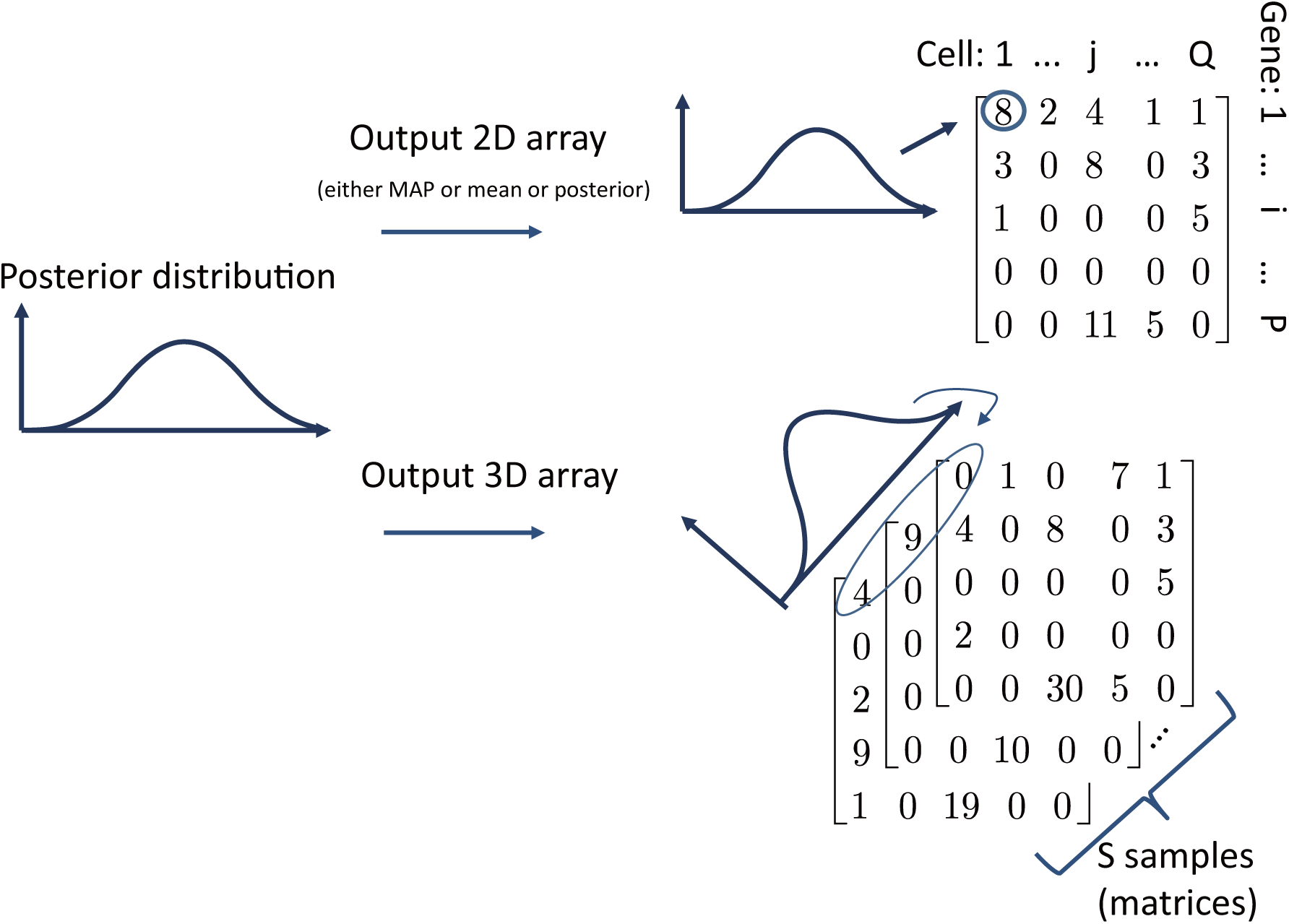
Output of bayNorm. For each gene in each cell, we have a posterior distribution as bayNorm is a Baysian method (See methods). Final bayNorm output is either S samples randomly sampled from the posterior distributions (3D arrays), or the mode or mean of the posterior used as point estimates (2D arrays).

**Supplementary Figure 2:**
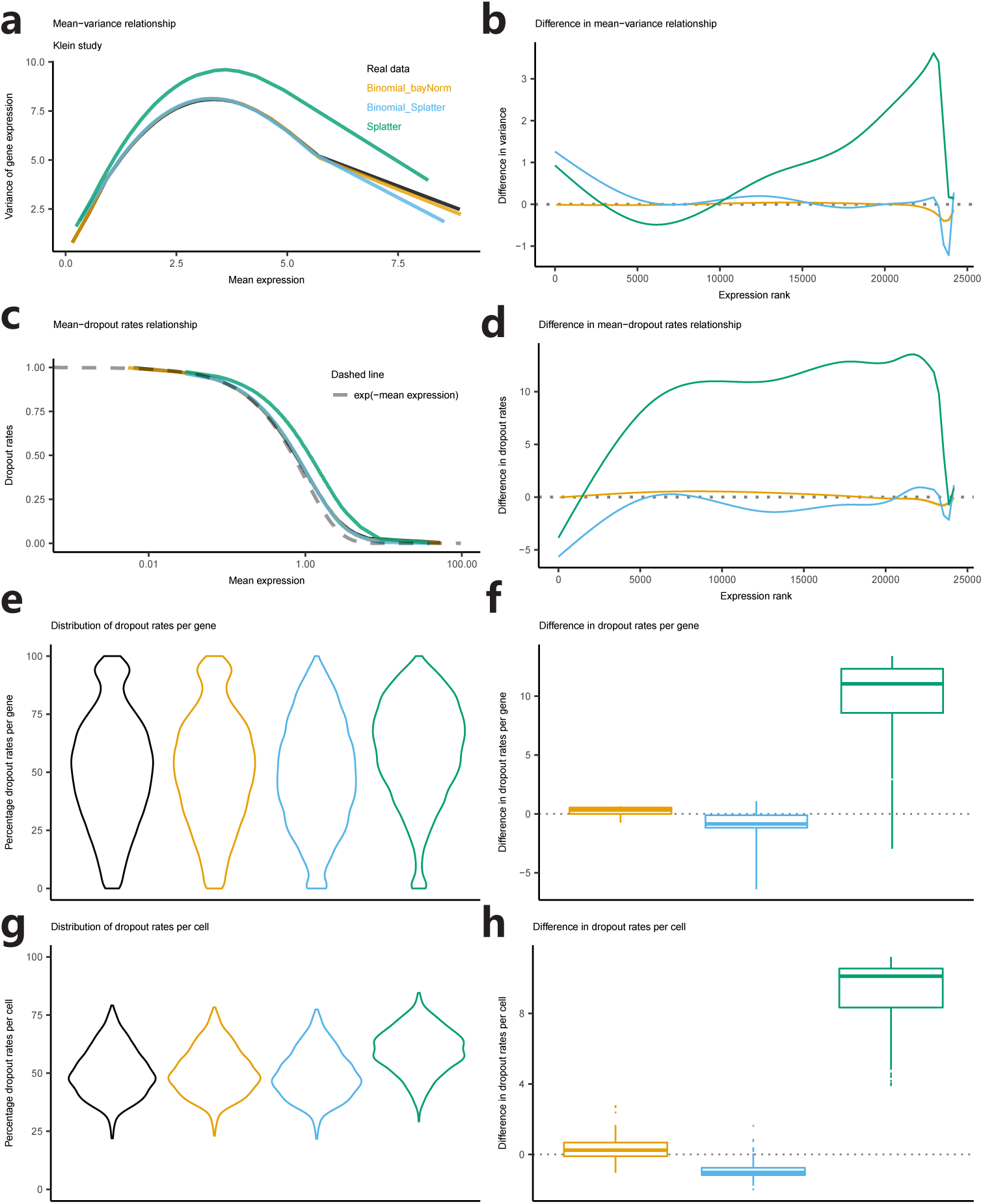
Simulation analysis based on the Klein study. Comparison between simulated data and experimental data in terms of: (a-b) variance-mean relationship, (c-d) dropout rates-mean relationship, (e-f) distribution of proportions of zeros per gene, (g-h) distribution of proportions zeros per cell. (b), (d), (f) and (h): ranked difference between statistics of experimental data and simulated data for (a), (c), (e) and (g) respectively. The smoothed lines in (a) and (c) were obtained by binning x values and calculating the mean of y values in each bin[1]. The smoothed lines in (b) and (d) were generated by ggplot2 (geom smooth with method set to “auto”).

**Supplementary Figure 3:**
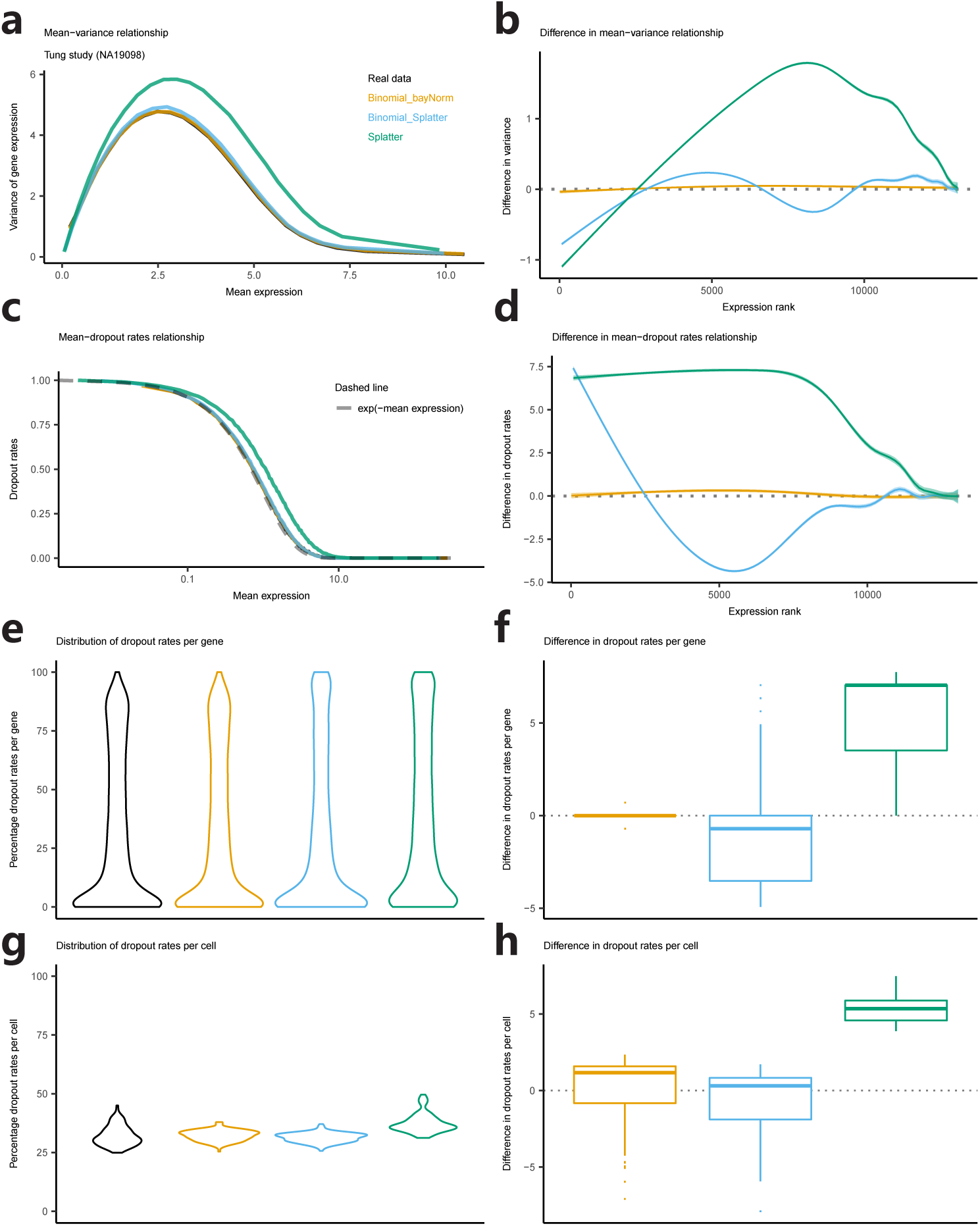
Simulation analysis based on the Tung study (Individual NA19098). Comparison between simulated data and real data in terms of: (a-b) variance-mean relationship, (c-d) dropout rates-mean relationship, (e-f) distribution of proportions of zeros per gene, (g-h) distribution of proportions zeros per cell. (b), (d), (f) and (h): ranked difference between statistics of experimental data and simulated data for (a), (c), (e) and (g) respectively. The smoothed lines in (a) and (c) were obtained by binning x values and calculating the mean of y values in each bin[1]. The smoothed lines in (b) and (d) were generated by ggplot2 (geom smooth with method set to “auto”).

**Supplementary Figure 4:**
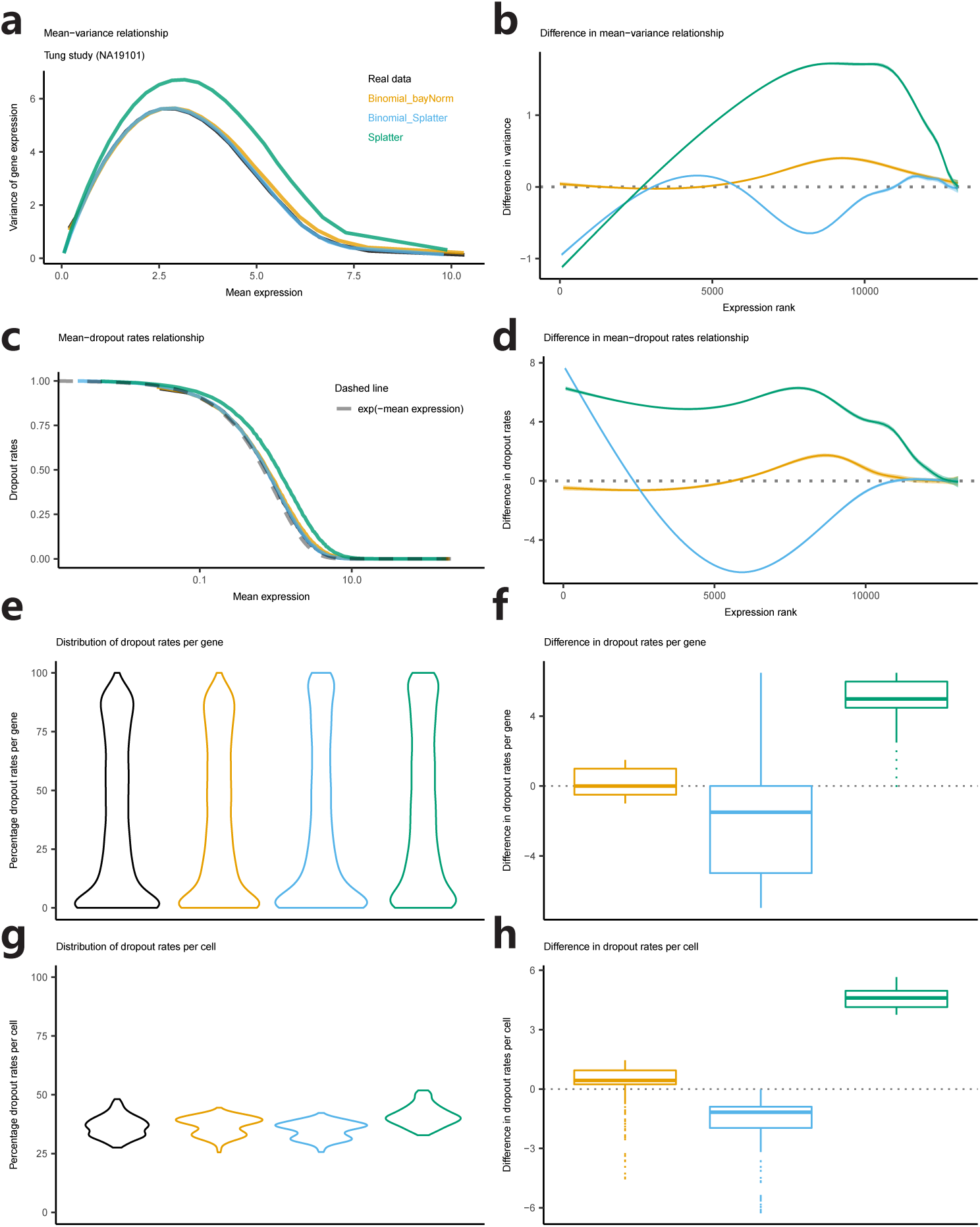
Simulation analysis based on the Tung study (Individual NA19101). Comparison between simulated data and real data in terms of: (a-b) variance-mean relationship, (c-d) dropout rates-mean relationship, (e-f) distribution of proportions of zeros per gene, (g-h) distribution of proportions zeros per cell. (b), (d), (f) and (h): ranked difference between statistics of experimental data and simulated data for (a), (c), (e) and (g) respectively. The smoothed lines in (a) and (c) were obtained by binning x values and calculating the mean of y values in each bin[1]. The smoothed lines in (b) and (d) were generated by ggplot2 (geom smooth with method set to “auto”).

**Supplementary Figure 5:**
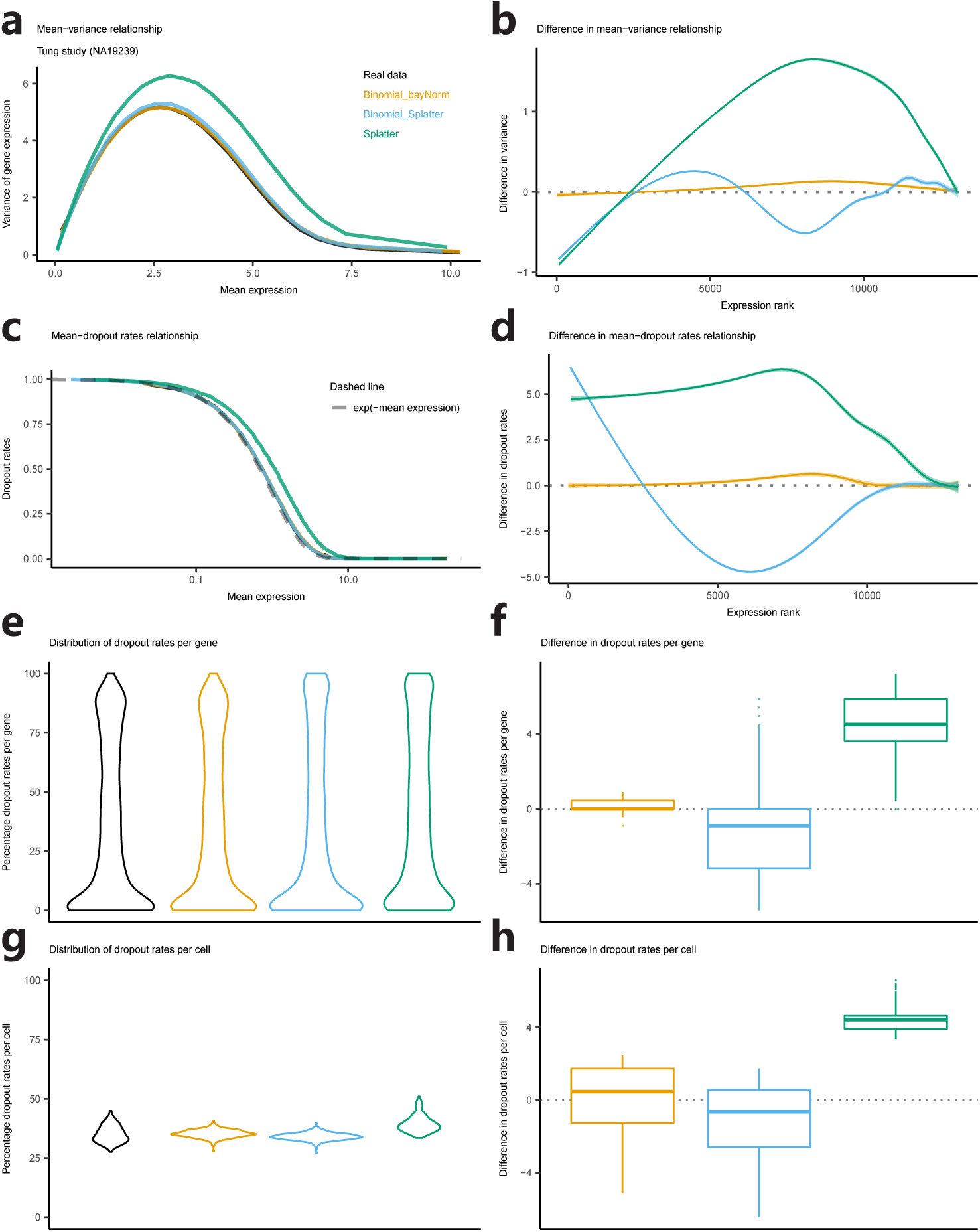
Simulation analysis based on the Tung study (Individual NA19239). Comparison between simulated data and real data in terms of: (a-b) variance-mean relationship, (c-d) dropout rates-Mean relationship, (e-f) distribution of proportions of zeros per gene, (g-h) distribution of proportions zeros per cell. (b), (d), (f) and (h): ranked difference between statistics of experimental data and simulated data for (a), (c), (e) and (g) respectively. The smoothed lines in (a) and (c) were obtained by binning x values and calculating the mean of y values in each bin[1]. The smoothed lines in (b) and (d) were generated by ggplot2 (geom smooth with method set to “auto”).

**Supplementary Figure 6:**
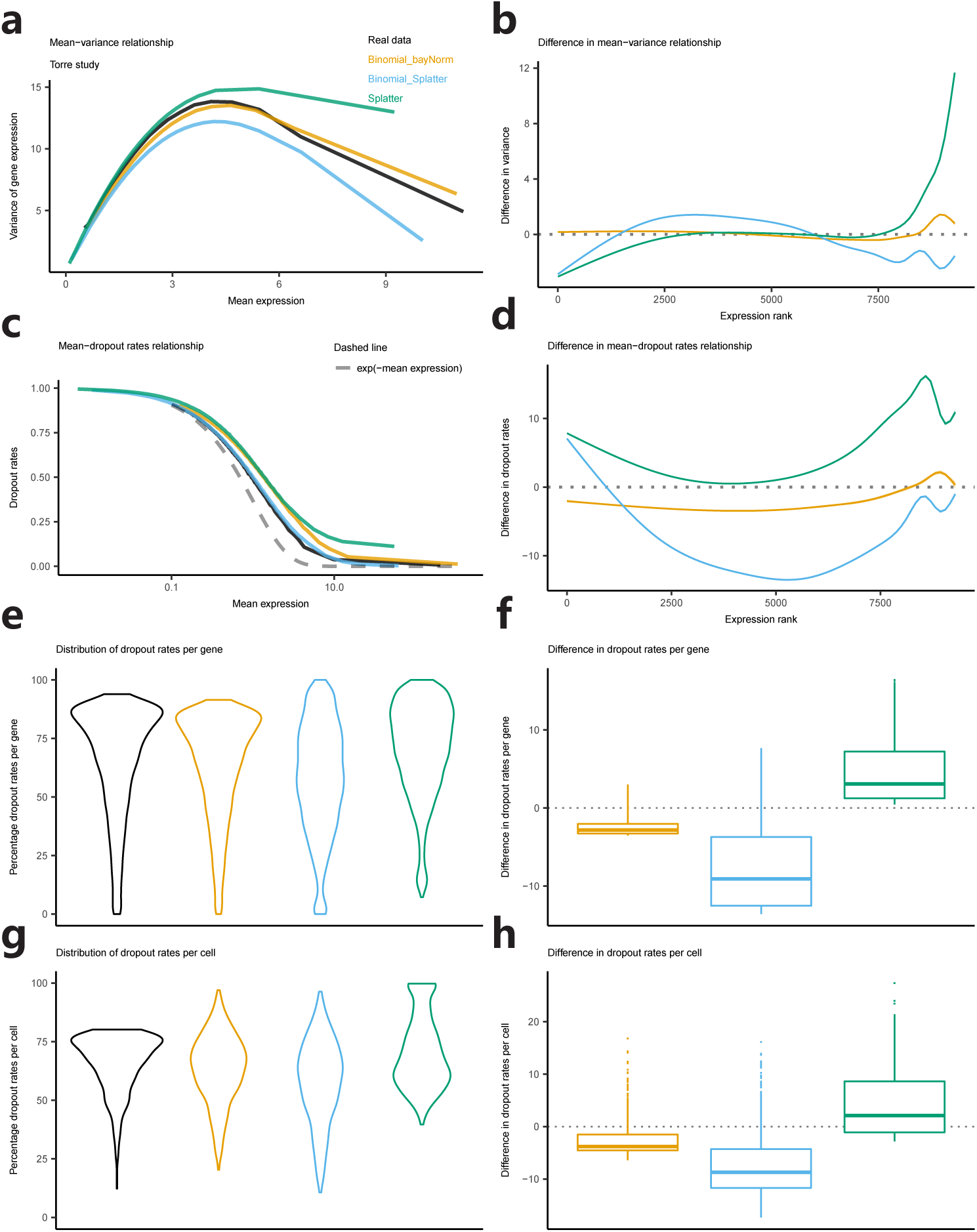
Simulation analysis based on the Torre study. Comparison between simulated data and real data in terms of: (a-b) variance-mean relationship, (c-d) dropout rates-mean relationship, (e-f) distribution of proportions of zeros per gene, (g-h) distribution of proportions zeros per cell. (b), (d), (f) and (h): ranked difference between statistics of experimental data and simulated data for (a), (c), (e) and (g) respectively. The smoothed lines in (a) and (c) were obtained by binning x values and calculating the mean of y values in each bin[1]. The smoothed lines in (b) and (d) were generated by ggplot2 (geom smooth with method set to “auto”).

**Supplementary Figure 7:**
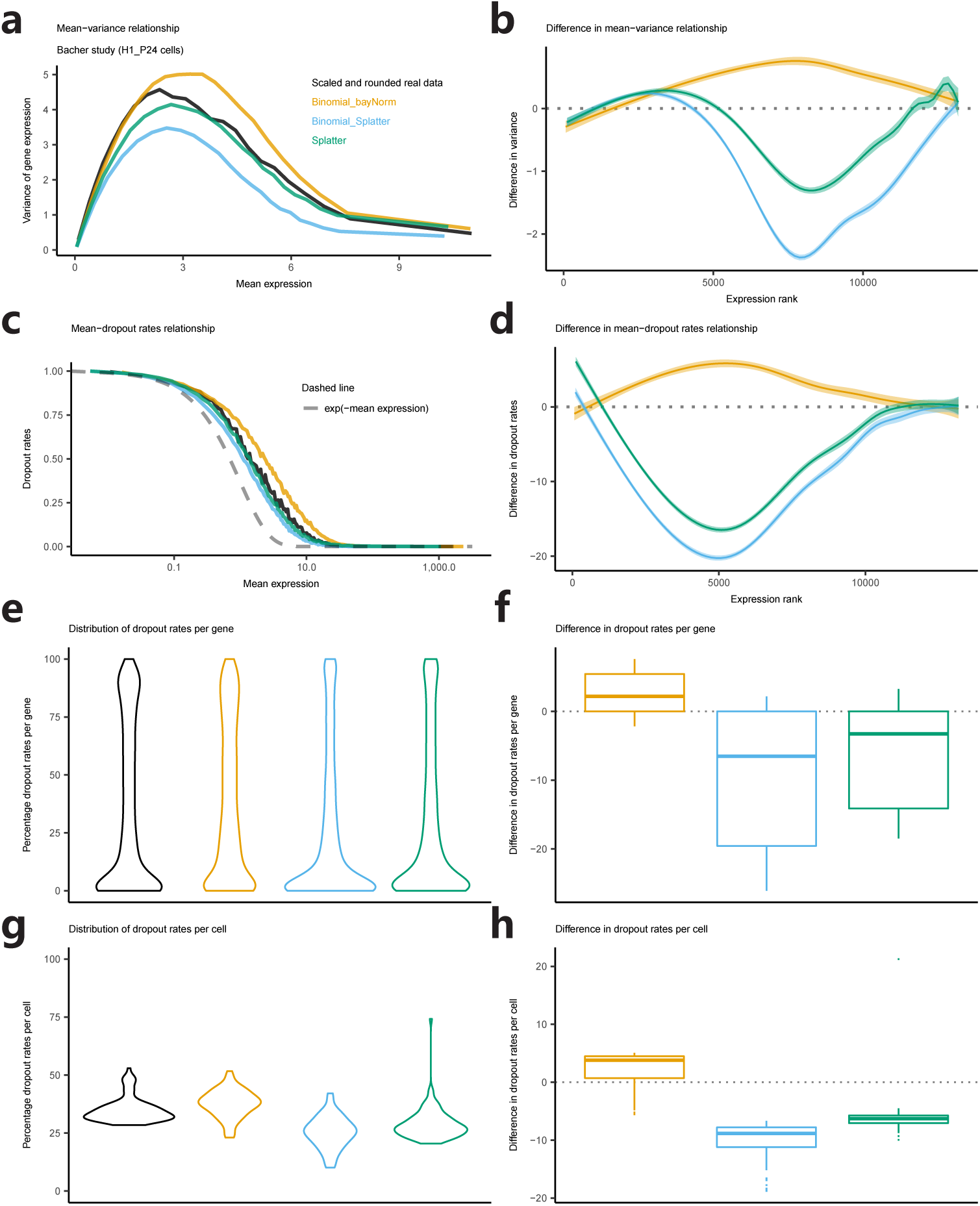
SSimulation analysis based on H1_P24 cells in the Bacher study. Comparison between simulated data and real data in terms of: (a-b) variance-mean relationship, (c-d) dropout rates-mean relationship, (e-f) distribution of proportions of zeros per gene, (g-h) distribution of proportions zeros per cell. (b), (d), (f) and (h): ranked difference between statistics of experimental data and simulated data for (a), (c), (e) and (g) respectively. The smoothed lines in (a) and (c) were obtained by binning x values and calculating the mean of y values in each bin33. The smoothed lines in (b) and (d) were generated by ggplot2 (geom smooth with method set to “auto”). Experimental data were scaled by 20 and rounded before being used as input of the three simulation protocols.

**Supplementary Figure 8:**
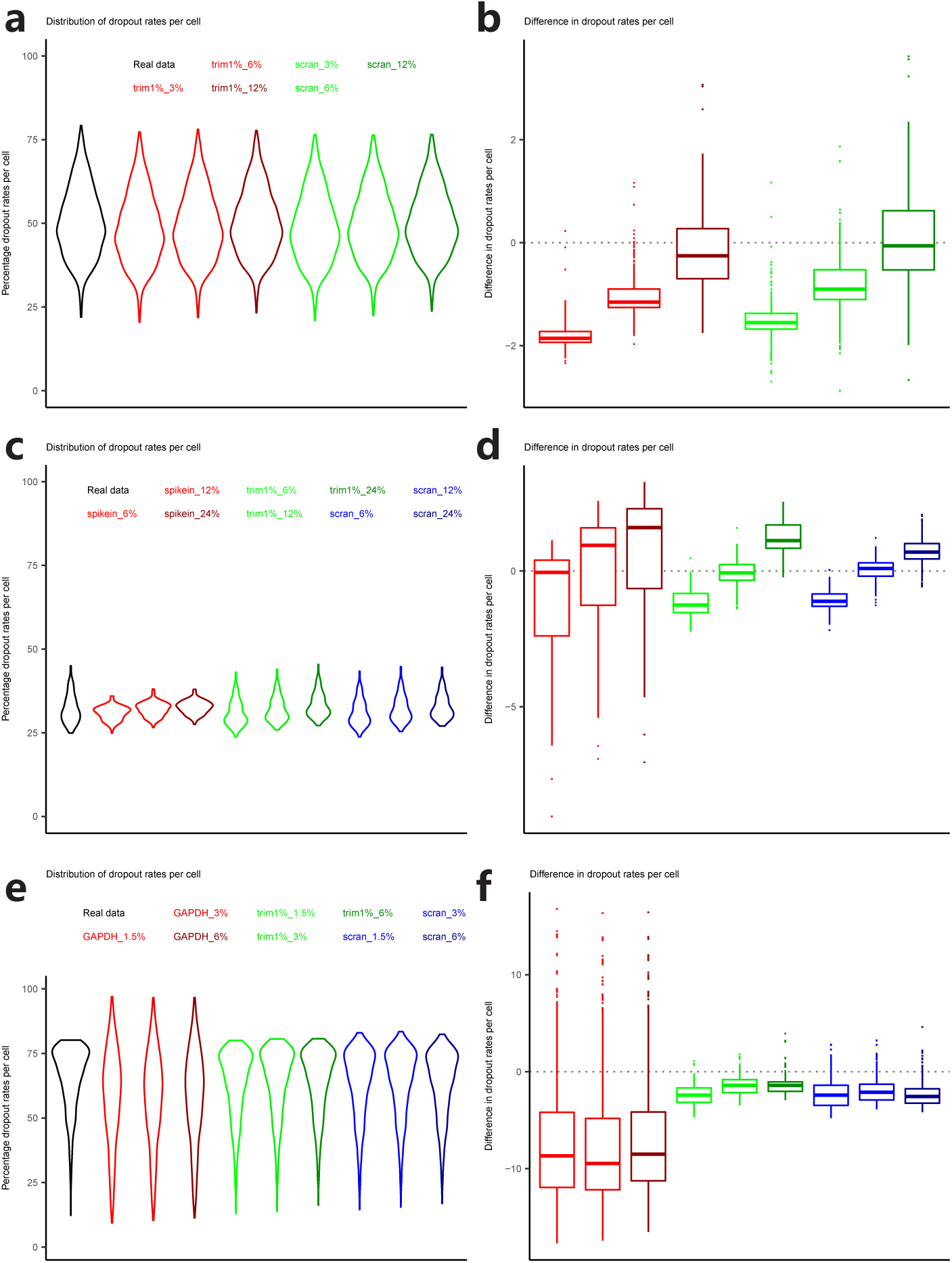
Impact of different mean capture efficiencies and different size factor estimates on the Binomial Splatter simulation protocol. Results are based on (a-b) the Klein study. (c-d) the NA19098 sample from the Tung study. (e-f) the Torre study. Scaling factors were estimated using different methods: (1) trim1%: 1% of counts were trimmed from each end of the counts in a specific cell before computing the mean. (2): “scran”: scaling factors were estimated with the R package scran[2]. (3) “spikein”: total counts of observed spike-ins in each cell were used as scaling factors. (4) “GAPDH”: the expression of the housekeeping gene GAPDH was used as scaling factors. The percentage at the end of each label indicates the mean capture efficiency *< β >* (see Methods).

**Supplementary Figure 9:**
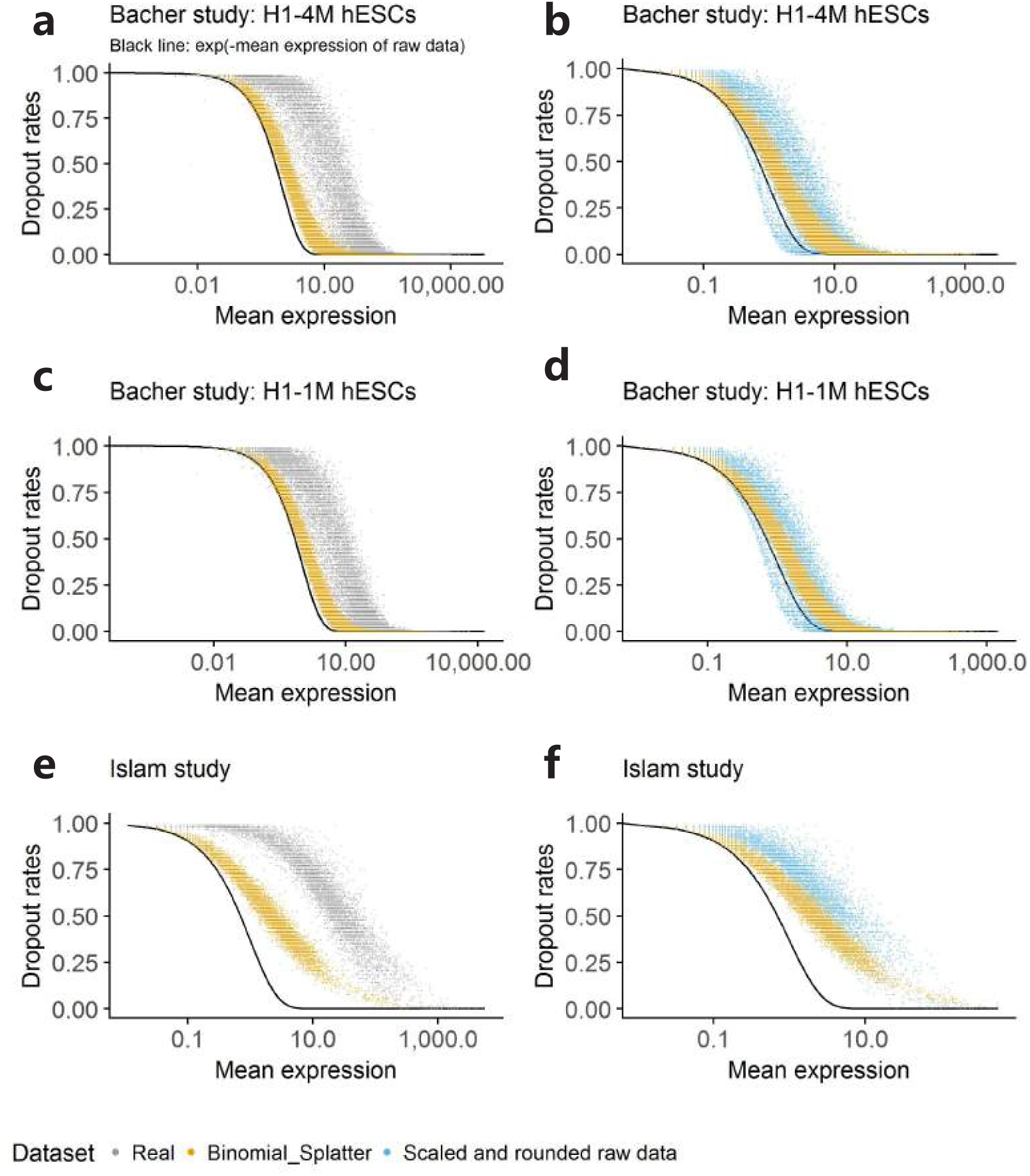
Comparison between simulated data and raw experimental data in terms of the relationship between dropout rates and mean expression. (a-b) non-UMI data from the Bacher study (H1 hESCs from the 4 million mapped reads group). In (b) raw experimental data were divided by 20 and rounded. (c-d) non-UMI data from the Bacher study (H1 hESCs from the 1 million mapped reads group). In (d) raw experimental data were divided by 10 and rounded. (e-f) non-UMI data from the Islam study. In (f) raw experimental data were divided by 10 and rounded. (b), (d) and (f) are comparisons between Binomial Splatter simulated data and scaled and rounded real experimental data. Parameters in Binomial Splatter simulations were generated from scaled raw data as illustrated in the Supplementary Note 4.

**Supplementary Figure 10:**
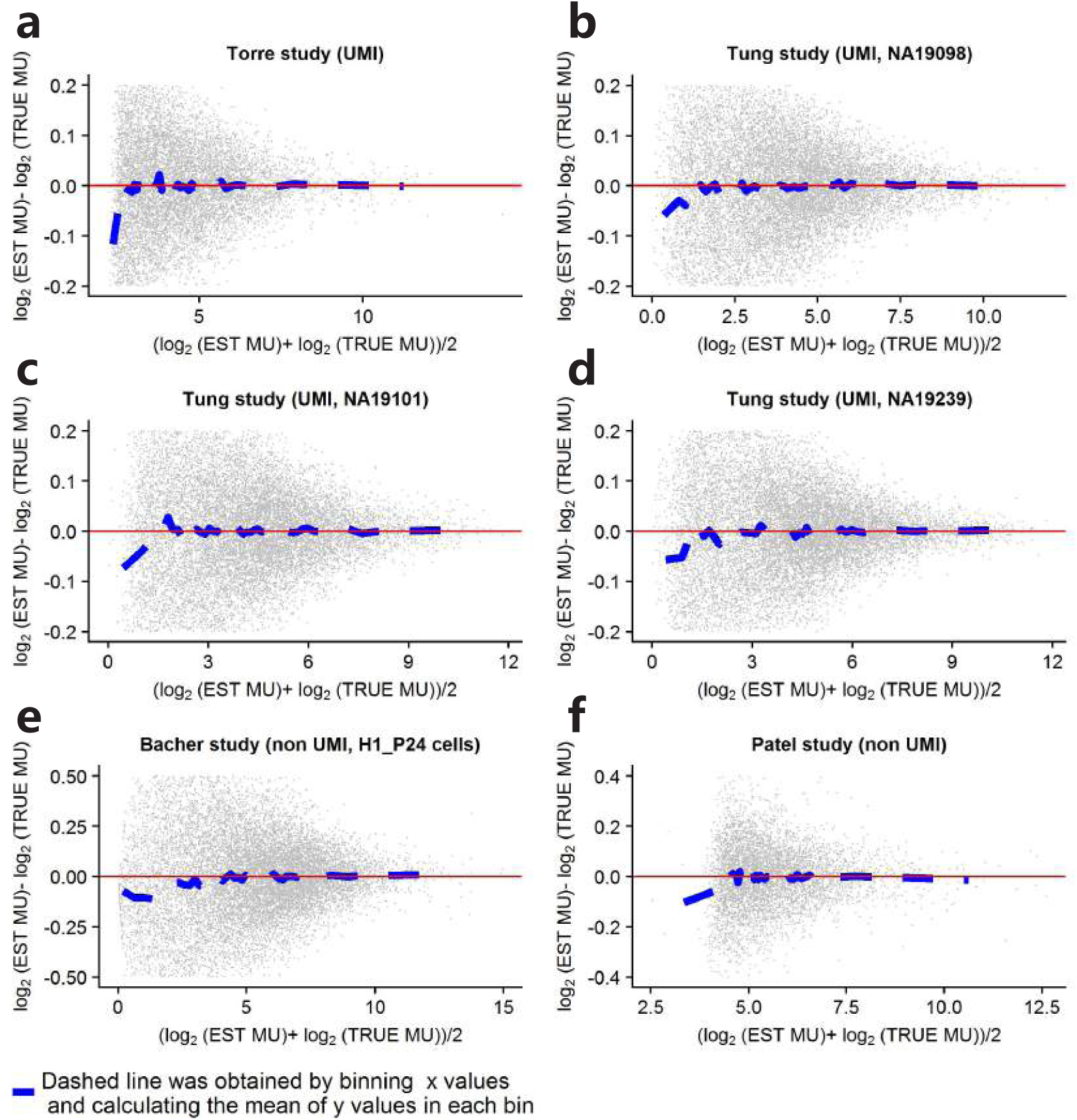
MA plots based on simulated data using the Binomial bayNorm protocol. TRUE MU stands for the MME estimated *µ* output from bayNorm and “EST MU” stands for the mean expression of Binomial bayNorm simulated data scaled by the *β* used in bayNorm. Binomial bayNorm simulation protocol was applied to (a) the Torre study, (b) individual NA19098 from the Tung study, (c) individual NA19101, (d) individual NA19239, (e) the Bacher study and (f) the Patel study. The dashed line was obtained by binning x values and calculating the mean of y values in each bin[1].

**Supplementary Figure 11:**
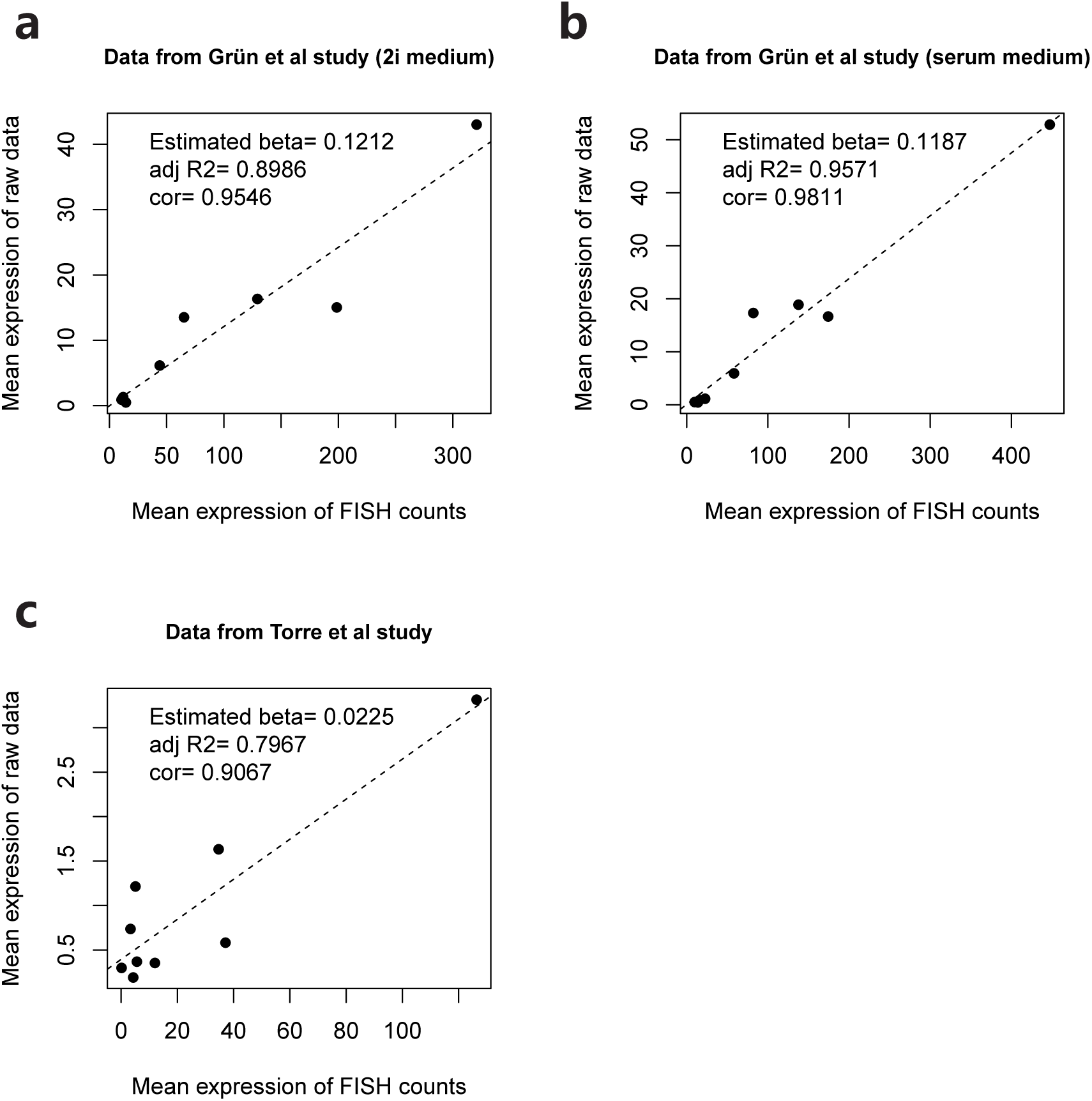
Linear regression of mean expression of scRNA-seq experimental raw data vs smFISH data. (a) 2i medium single cell data from the Grn study. (b) Serum medium single cell data from the Grn study. (c) Data from the Torre study. The coefficient of explanatory variable of linear regression is used as mean beta.

**Supplementary Figure 12:**
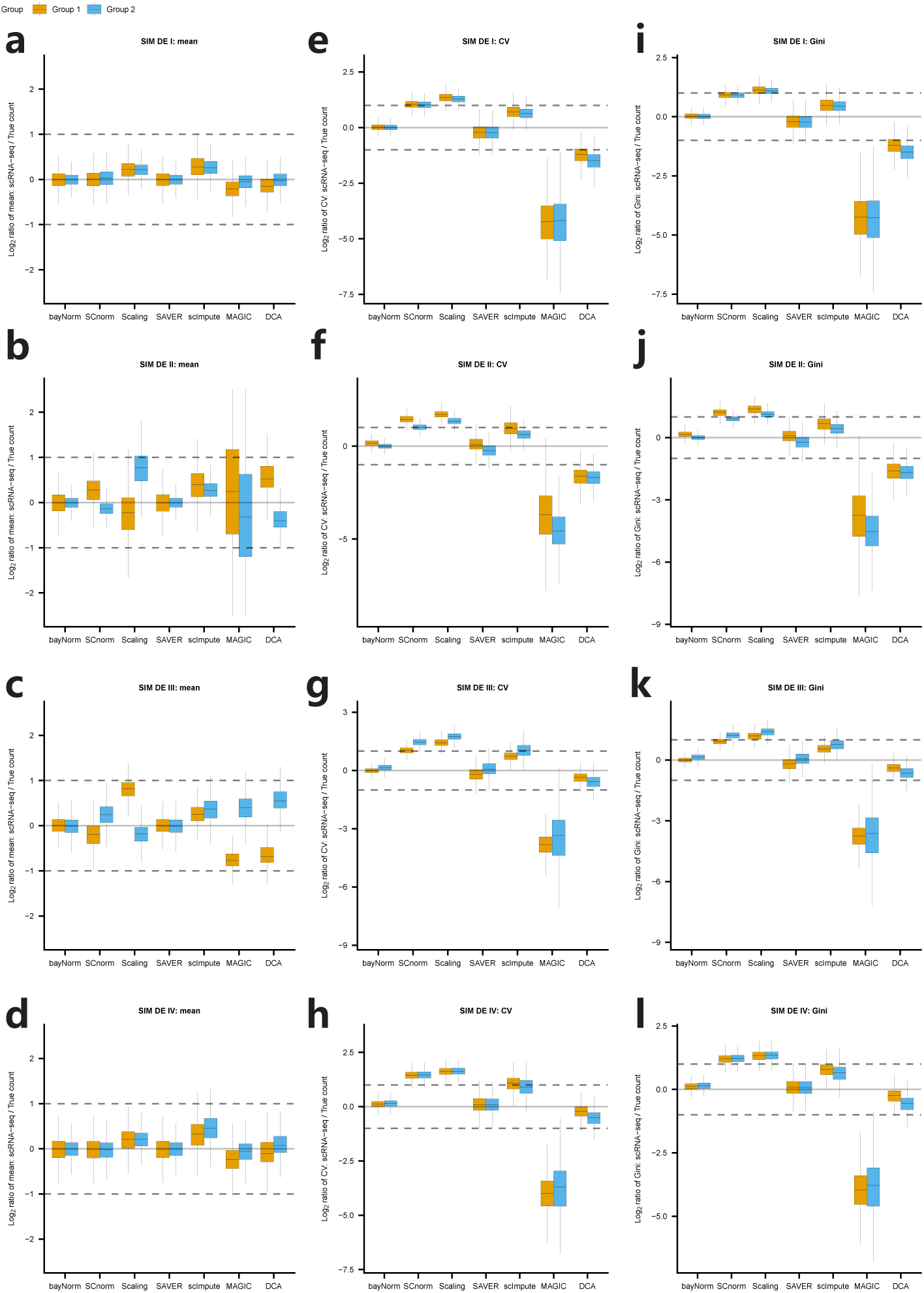
Recovering the mean, CV and Gini of gene expression using simulated scRNA-seq data. For the four simulation studies: SIM DE I-IV (See Supplementary note 1 and 2 for details about simulation studies), Log_2_ ratio between true simulated data (dataset before binomial downsampling) and normalized simulated scRNA-seq data for mean gene expression (a-d), CV (e-h) and Gini coefficients (i-l). (a), (e) and (i) are based on SIM DE I simulated data. (b), (f) and (j) are based on SIM DE II simulated data. (c), (g) and (k) are based on SIM DE III simulated data. (d), (h) and (l) are based on SIM DE IV simulated data. Except for the bayNorm and scaling methods, the normalized datasets have been divided by their corresponding mean capture efficiencies (either 0.1 or 0.05) for a fair comparison.

**Supplementary Figure 13:**
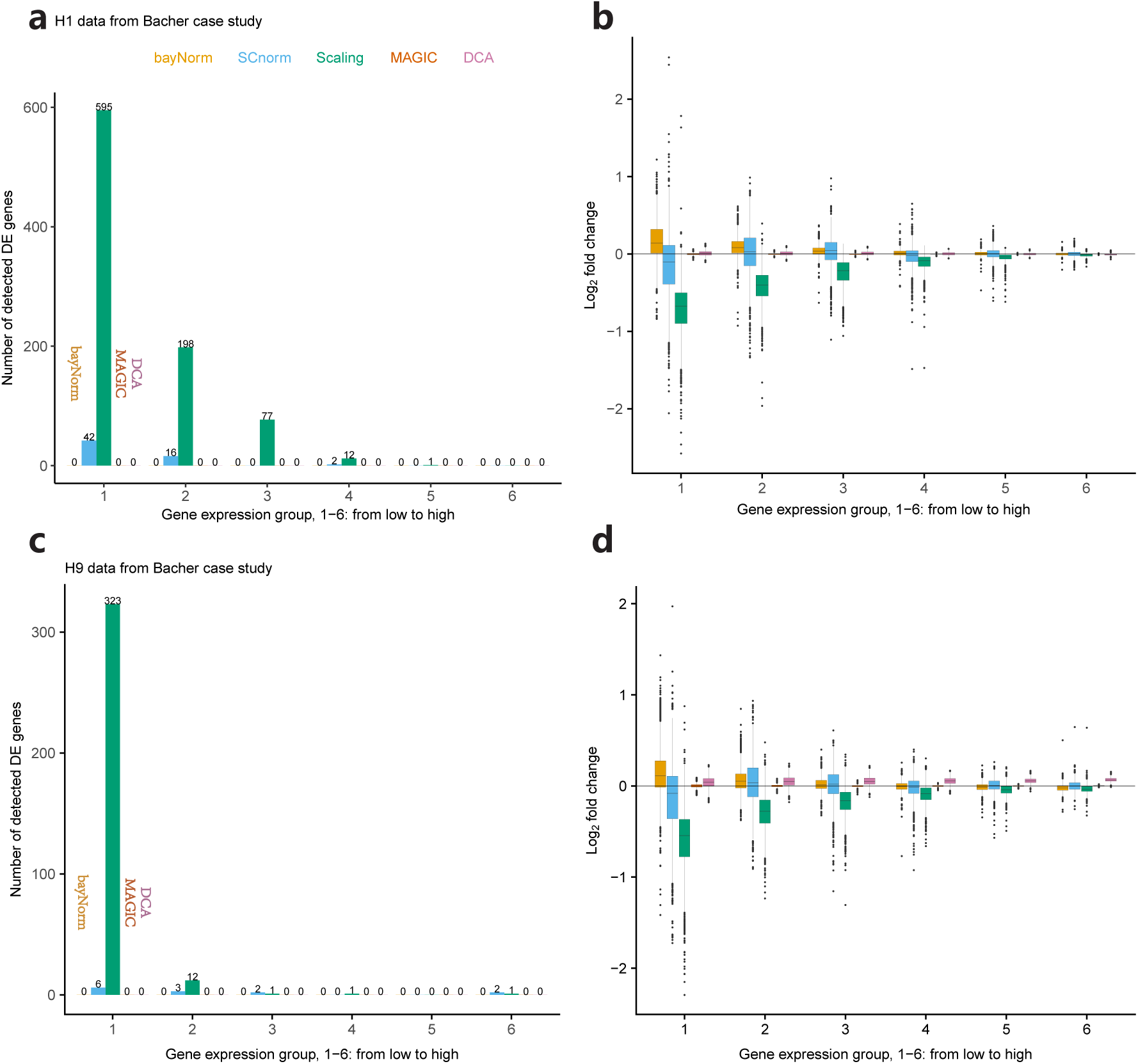
bayNorm correction for differences in sequencing depths. Data from H1 (a-b, 13181 genes in total) and H9 (c-d, 13195 genes in total) hESC cells are shown. (a) Number of DE genes called by MAST as a function of gene expression groups (*P*_MAST_ *<* 0.05). (b) Log_2_fold change as a function of gene expression group. In (a) bayNorm is based on 20 posterior samples (3D array). In (b), bayNorm is based on the mean of posteriors (2D array).

**Supplementary Figure 14:**
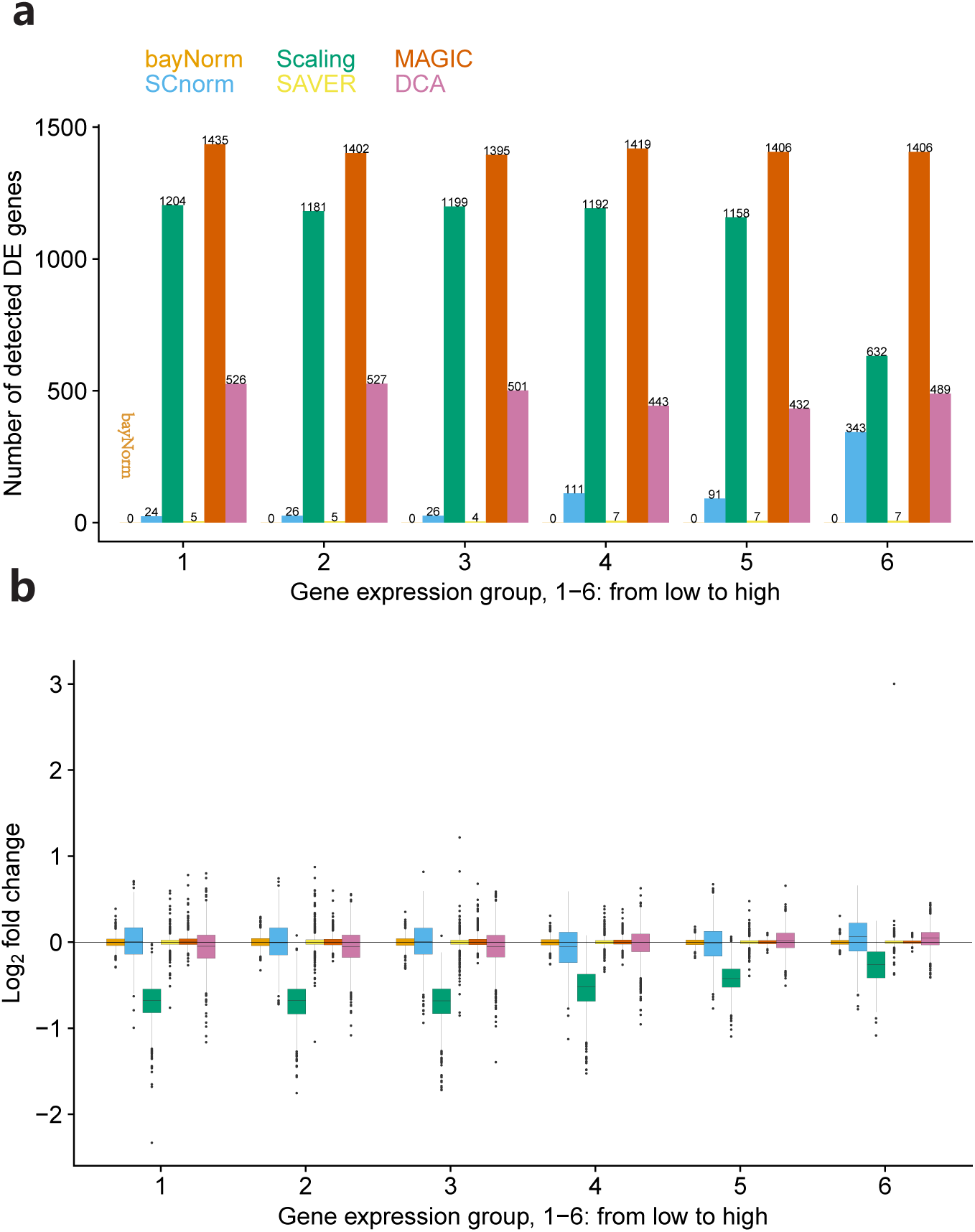
Simulation analysis, SIM noDE study I (See Supplementary note 1 and 2 for details about simulation studies). (a) Number of detected DE genes (MAST) as a function of expression group (*P*_MAST_ *<* 0.05, 9999 genes in total). (b) Log_2_fold change of mean expression between two groups for different expression groups. For bayNorm and SAVER, 10 samples were generated and the median of p-values across the 10 samples was used in (a). In (b), bayNorm and SAVER are based on mean of posterior distributions.

**Supplementary Figure 15:**
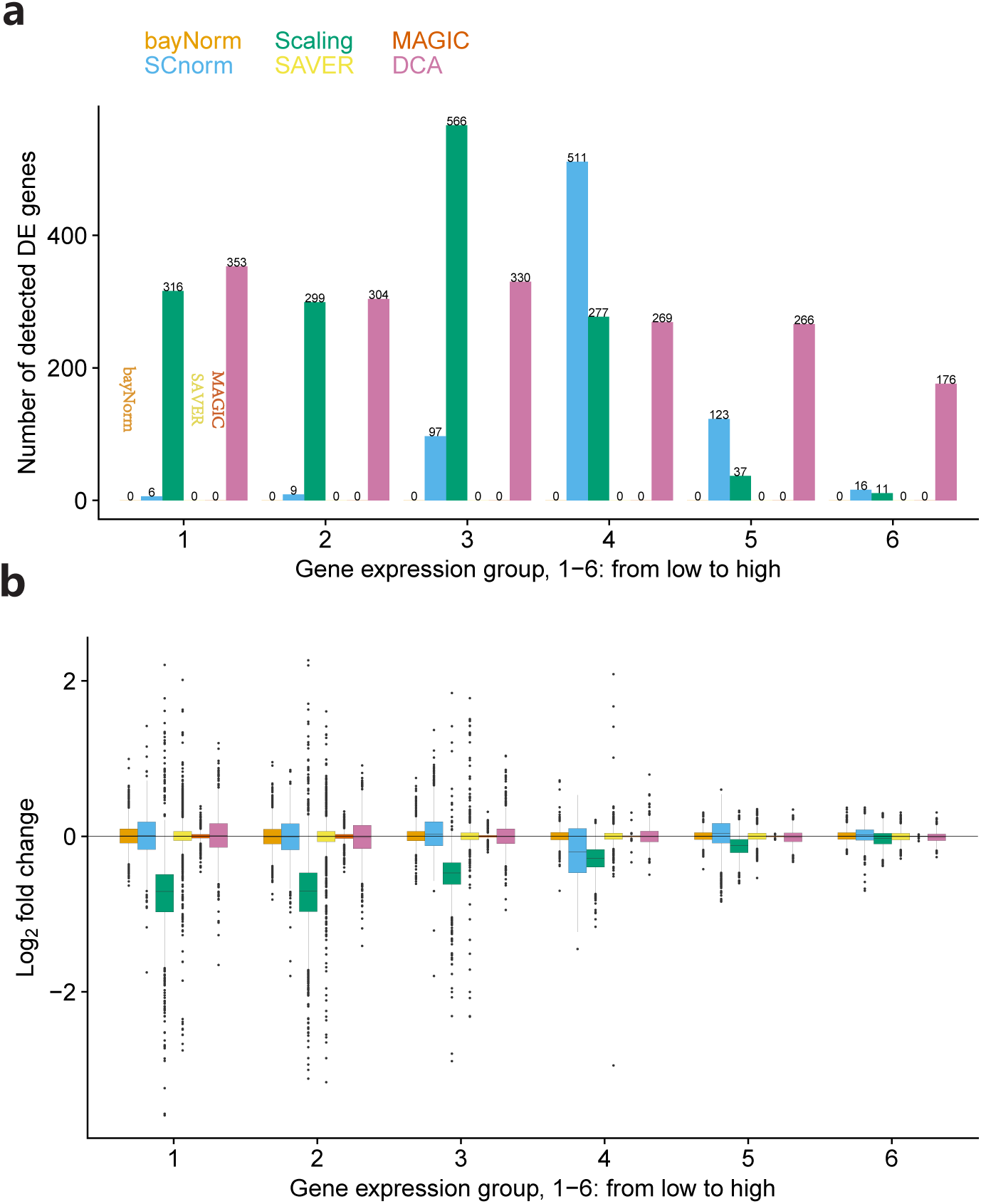
Simulation analysis, SIM noDE study II (See Supplementary note 1 and 2 for details about simulation studies). (a) Number of detected DE genes (MAST) as a function of expression group (*P*_MAST_ *<* 0.05, 9598 genes in total). (b) Log_2_ fold change of mean expression between two groups for different expression groups. For bayNorm and SAVER, 10 samples were generated and the median of p-values across the 10 samples was used in (a). In (b), bayNorm and SAVER are based on mean of posterior distributions.

**Supplementary Figure 16:**
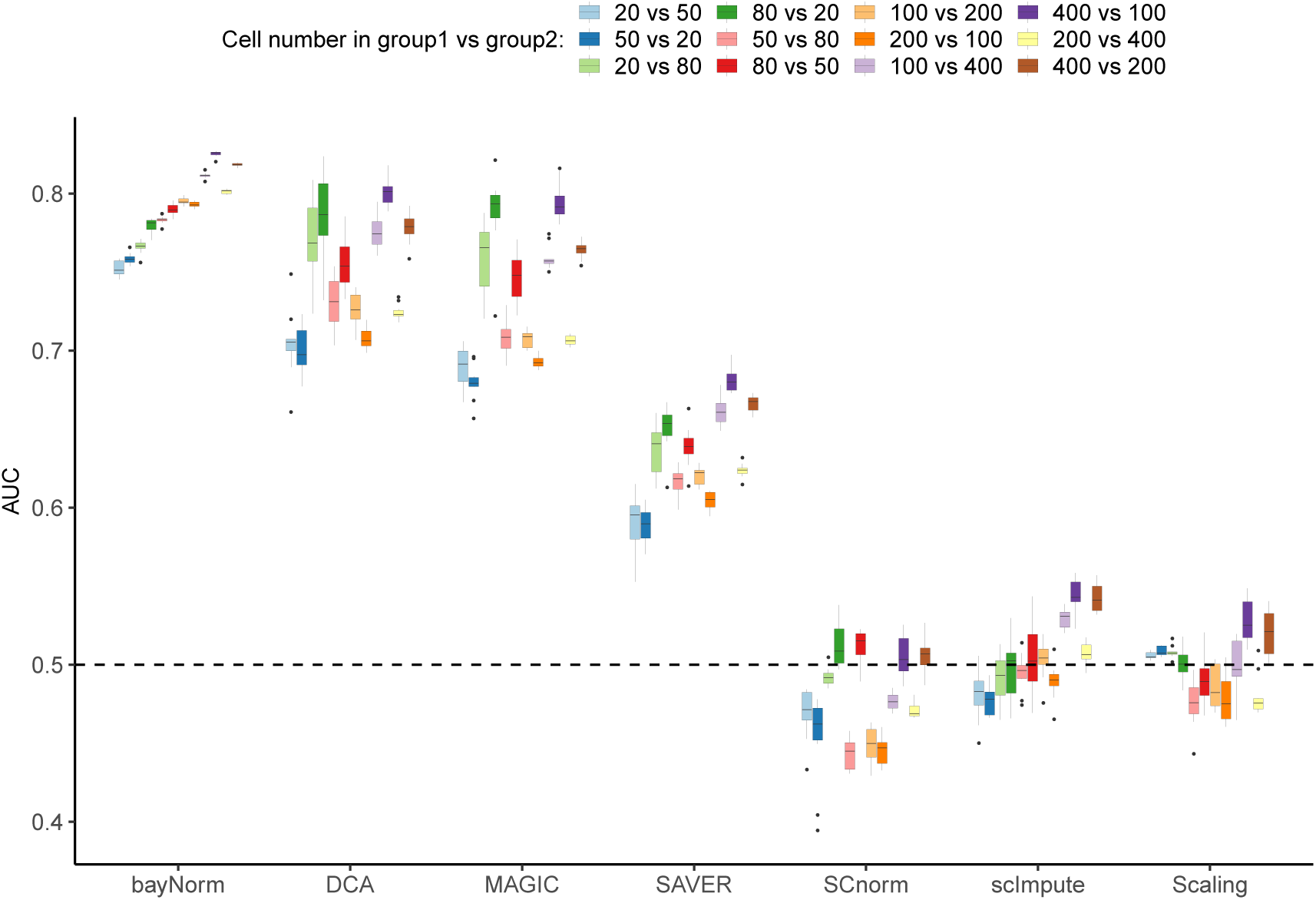
DE detection for unbalanced groups of cells (UMI data from the Soumillon study). Ten samples of 20, 50, 80, 100, 200 and 400 cells were randomly selected from each group. DE detection was performed using MAST between groups as described at the top of the figure using a list of DE genes obtained from matched bulk RNA-seq data as a benchmark (1000 genes with the largest magnitude of log fold-change between the D3T0 and D3T7 samples)[3].

**Supplementary Figure 17:**
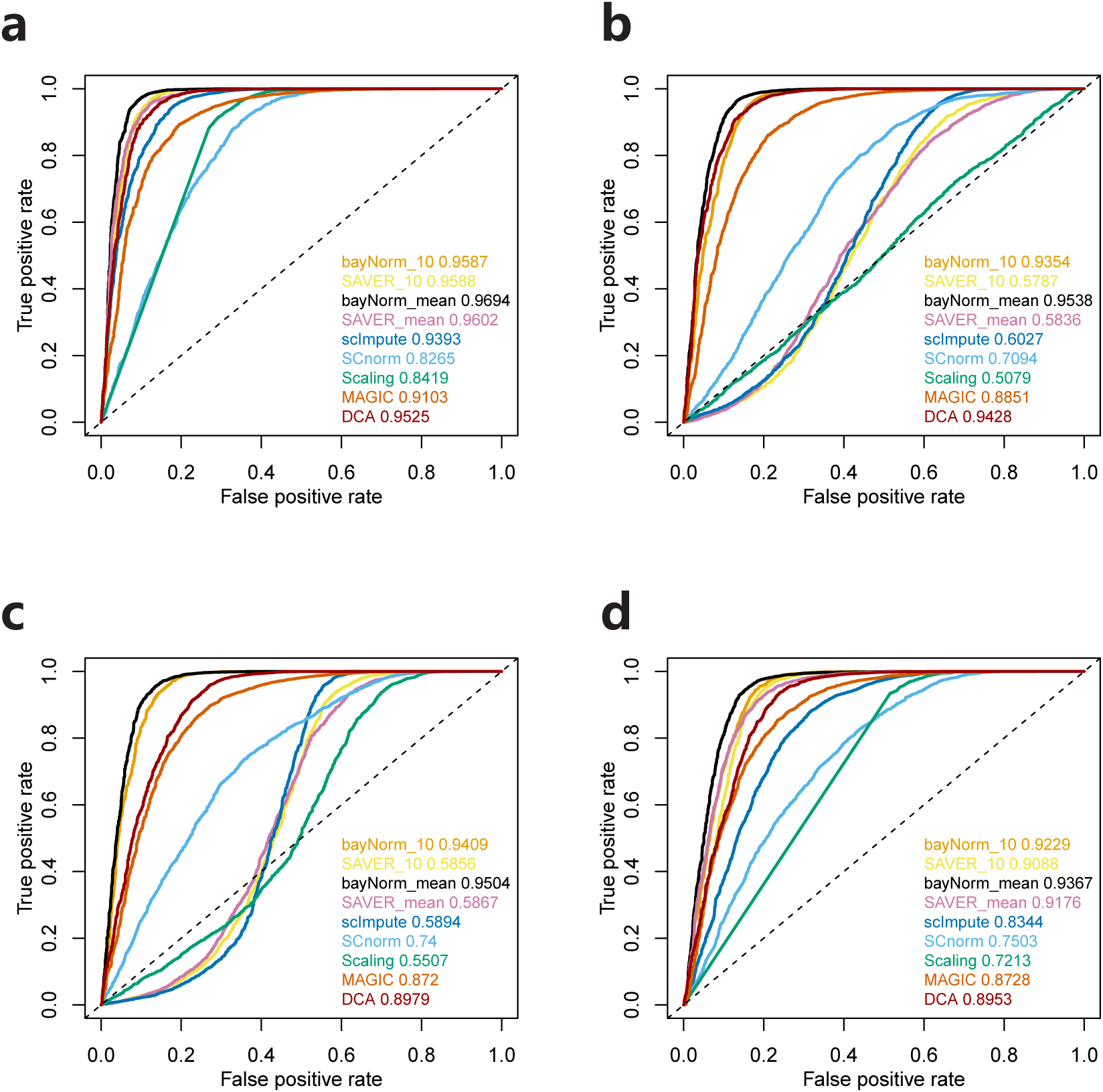
DE analysis on simulated scRNA-seq data, SIM DE study (See Supplementary note 1 and 2 for details about simulation studies). (a-d) represent four simulation scenarios and DE detection is based on MAST. (a) SIM I: mean capture efficiencies are set to 0.1 for the two groups. (b) SIM II: mean capture efficiencies are set to 0.05 and 0.1 in group 1 and group 2 respectively. (c) SIM III: mean capture efficiencies are set to 0.1 and 0.05 in group 1 and group 2 respectively. (d) SIM IV: mean capture efficiencies are set to 0.05 in both groups. 2000 out of 10000 genes were simulated to be DE genes in group 1. bayNorm 10 and SAVER 10 are based on 10 samples from posterior distributions (3D arrays). DE detection was performed on each sample and the median of adjusted MAST P-values were used.

**Supplementary Figure 18:**
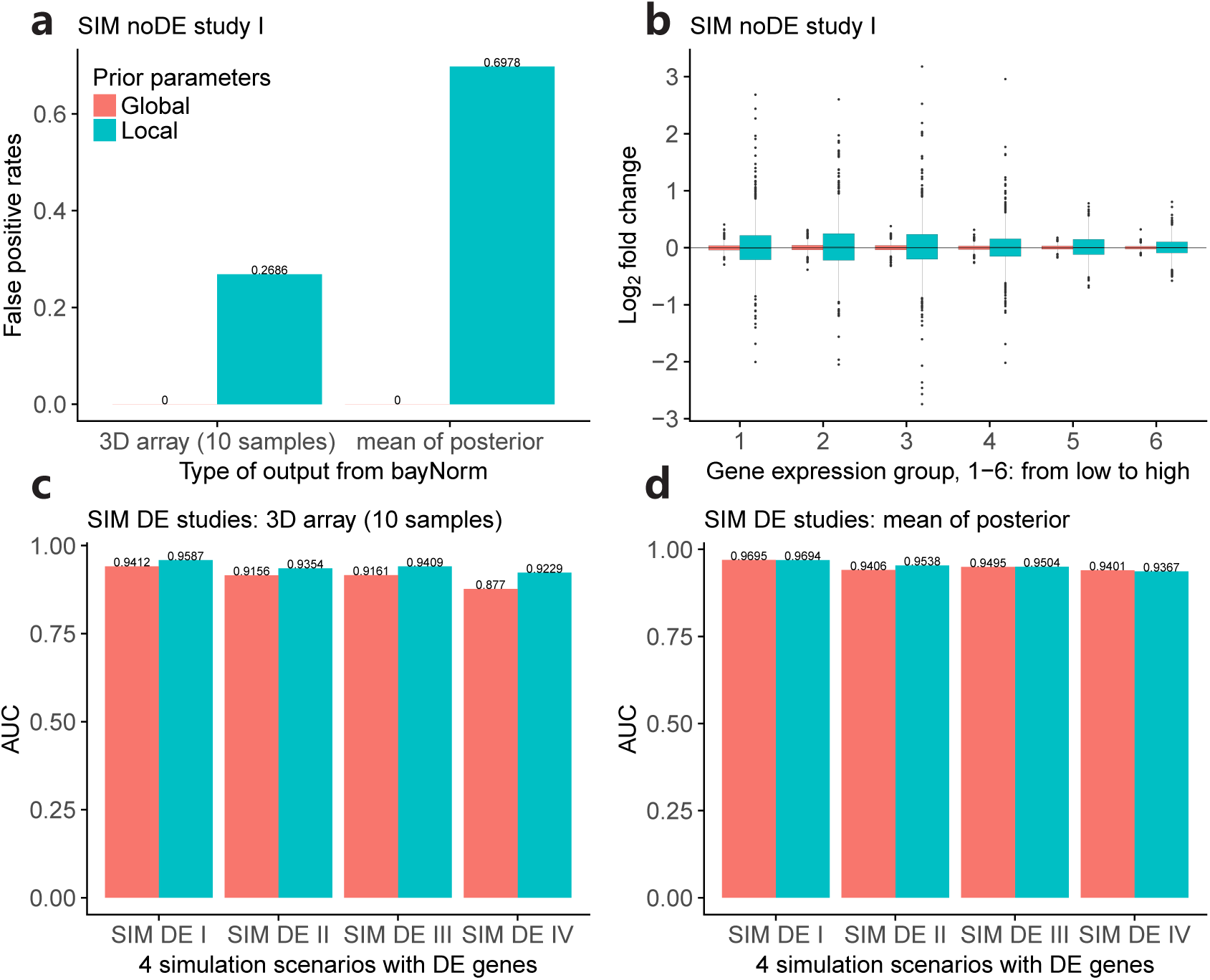
Impact of global or local priors on the DE detection for simulated samples with different sequencing depths. (a-b) Simulation analysis, SIM noDE study I, with no DE genes. (c-d) SIM DE studies I-IV where 2000 out of 10000 genes were simulated to be differentially expressed in the first group. (c) and (d) are based on 10 samples (3D array) and mean (2D array) output from bayNorm respectively.

**Supplementary Figure 19:**
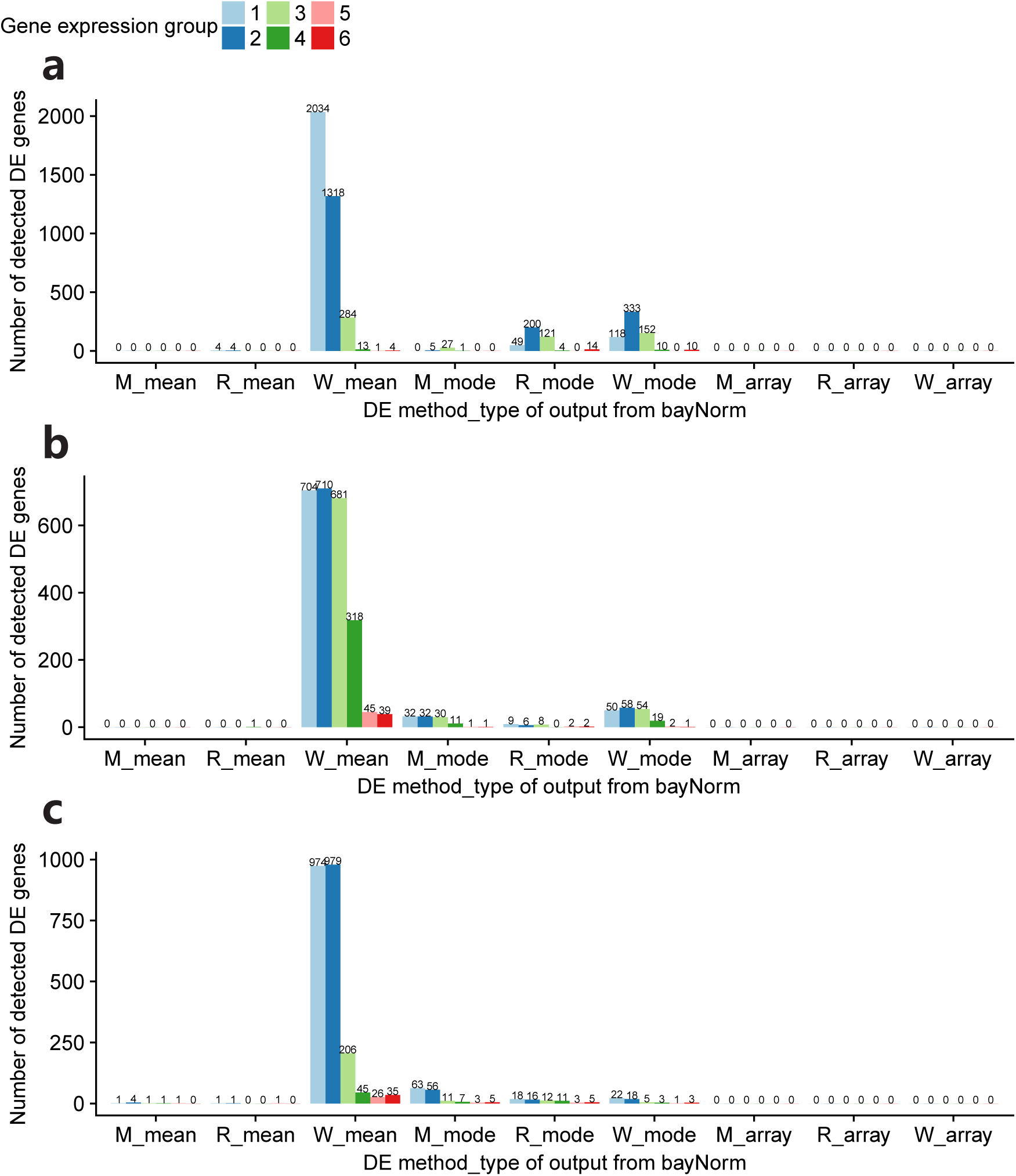
Impact of different DE methods and different types of output from bayNorm on samples with different sequencing depths. M stands for MAST, R stands for ROTS and W stands for Wilcoxon test. Mean (2D array), mode (2D array) and array (3D array) stands for the three different types of output from bayNorm (see Fig S1). For the 3D array output from bayNorm, each DE method was applied on each one of 10 samples from the posterior distribution, and the median of P-values was used. (a) H1 hESC data from the Bacher study. (b) Simulated data, SIM noDE study I. (c) Simulated data, SIM noDE study II. See Supplementary note 1 and 2 for details about simulation protocols. Genes were categorized into 6 groups according to their mean expression (1-low 6-high).

**Supplementary Figure 20:**
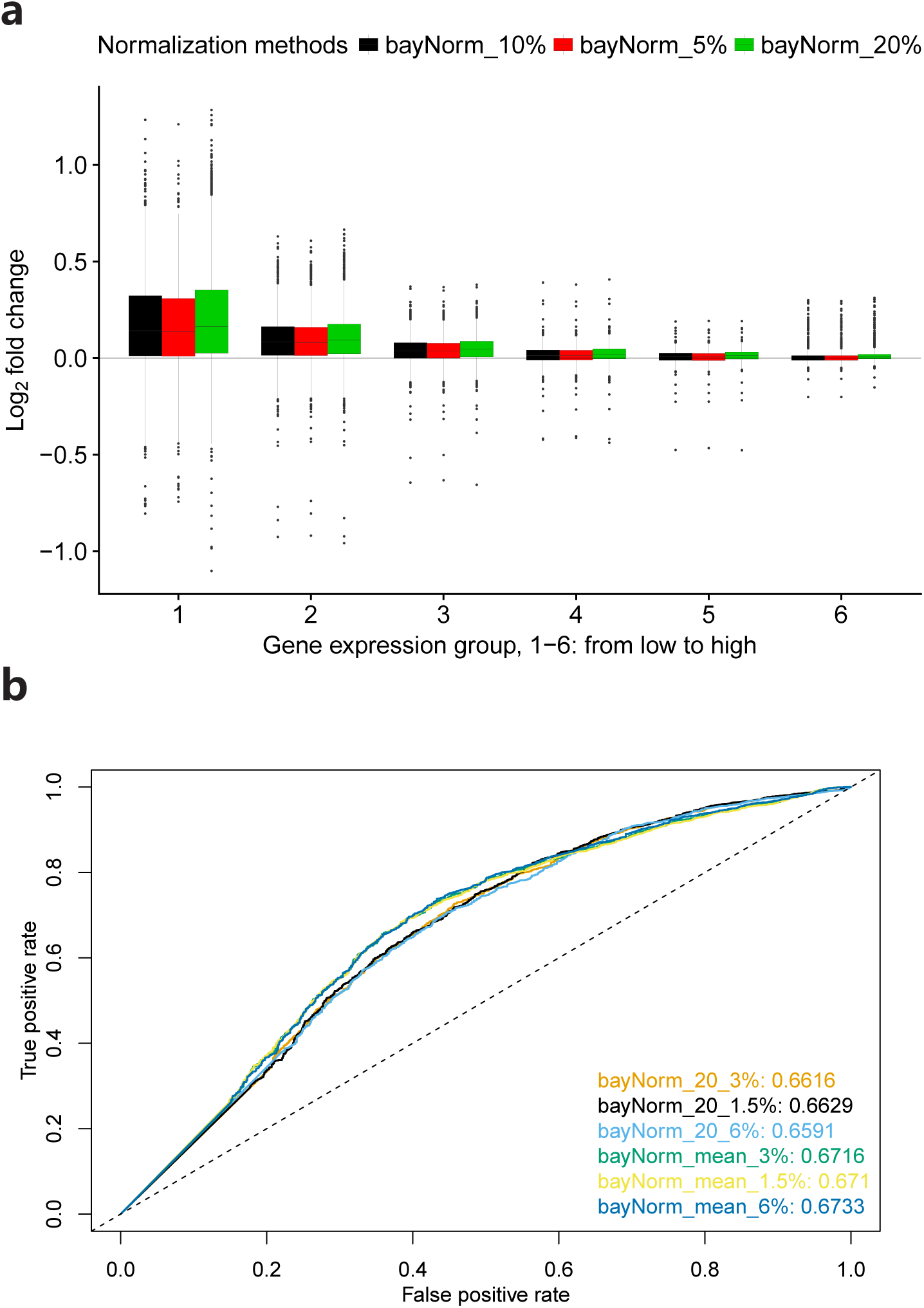
Impact of different mean capture efficiencies on DE analysis based on Bacher and Islam studies. (a) For data from the Bacher study, mean capture efficiencies were set to 5%, 10% or 20%. Results are based on mean of posterior output (2D array) from bayNorm. The result of DE detection was not shown as no genes were called DE at threshold 0.05. (b) Islam study. Mean capture efficiencies were set to 1.5%, 3% and 6%. mean (2D array) or 20 samples (3D array) were used for DE detection.

**Supplementary Figure 21:**
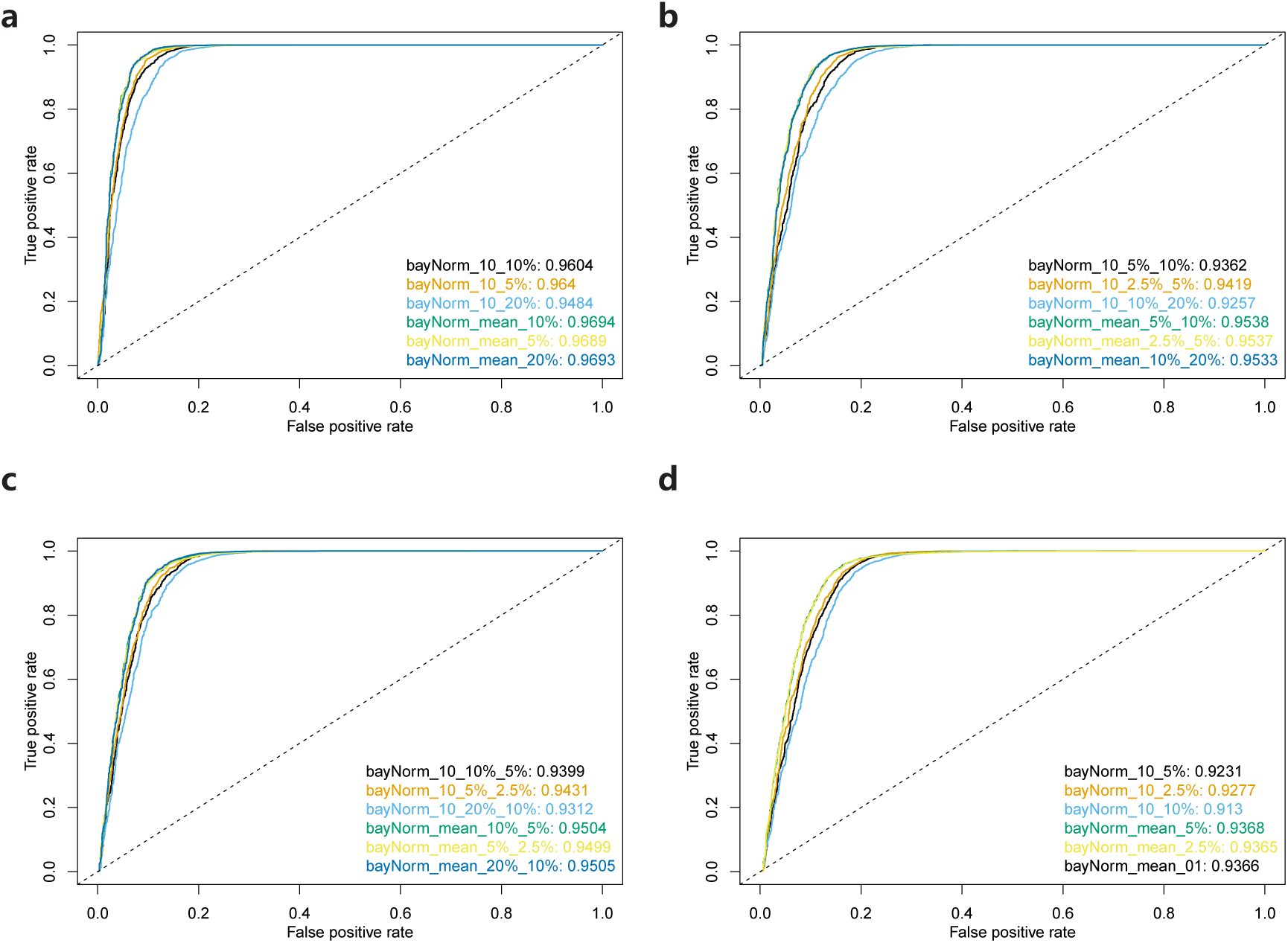
Impact of different mean capture efficiencies on DE detection in simulated studies (see Supplementary note 1 and 2). (a) SIM I from our SIM DE study. Mean capture efficiencies were set to 5%, 10% or 20% using either mean of posterior (2D array) or 10 samples generated from the posterior distributions (3D array) as normalized data. (b) SIM II from our SIM DE study. Mean capture efficiencies were set to twice or half of the original magnitude. (c) SIM III from our SIM DE study, Mean capture efficiencies were set to twice or half of the original magnitude. SIM IV from our SIM DE study. Mean capture efficiencies were set to 2.5%, 5% or 10%. mean stands for the mean versions output from bayNorm (2D array). Otherwise the number indicates the number of samples generated from posterior distribution (3D array). DE was performed on each sample, the median of MAST P-values were used.

**Supplementary Figure 22:**
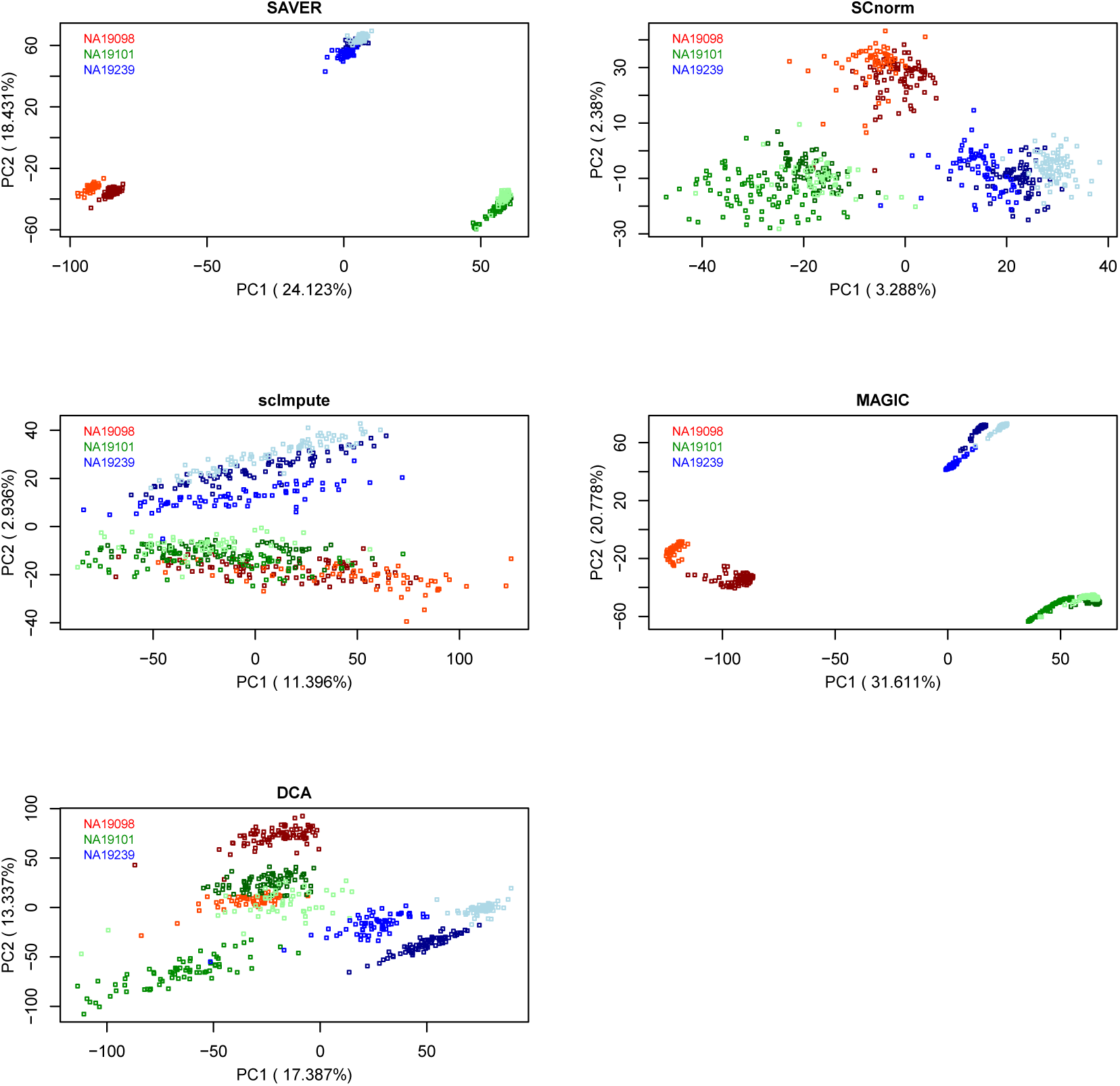
PCA plots of scRNA-seq data normalised using different methods (Tung study). PCA plot of SAVER normalized data is based on the mean versions output (2D array). Different colours represent different individuals. Different shades of the same colour stands for a specific batch within each individual.

**Supplementary Figure 23:**
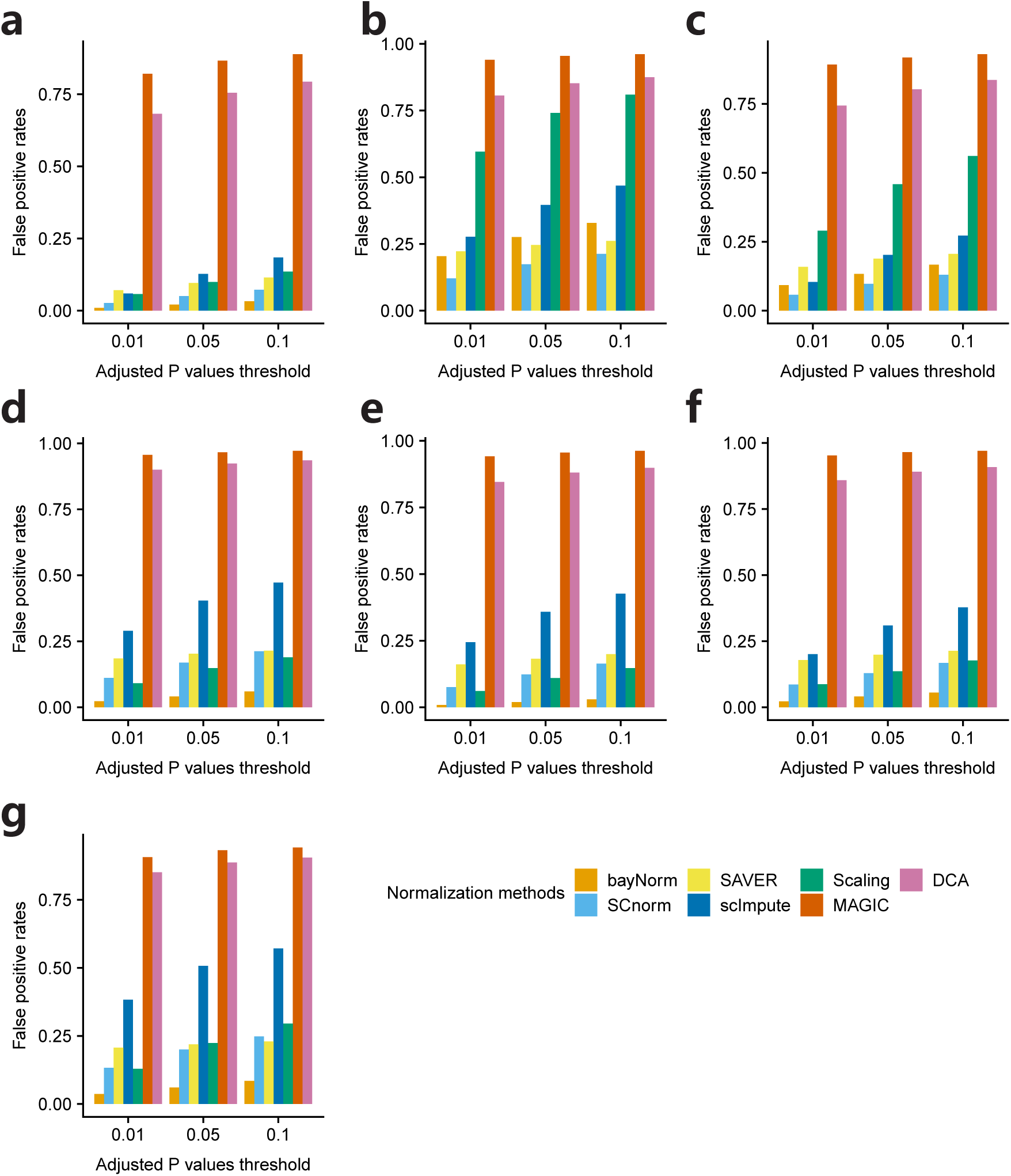
DE detection between scRNA-seq data for different batches within single individuals. (a-c) Individual NA19101, (d-f) Individual NA19239, (g) Individual NA19098. (a), (d) and (g) show DE detection between batch 1 and batch 3 (batch 2 was not considered as suggested in the Tung study). (b) and (e) show DE detection between batch 1 and batch 2. (c) and (f) show DE detection between batch 2 and batch 3. Results of bayNorm and SAVER are based on 5 samples from posterior distributions (3D array).

**Supplementary Figure 24:**
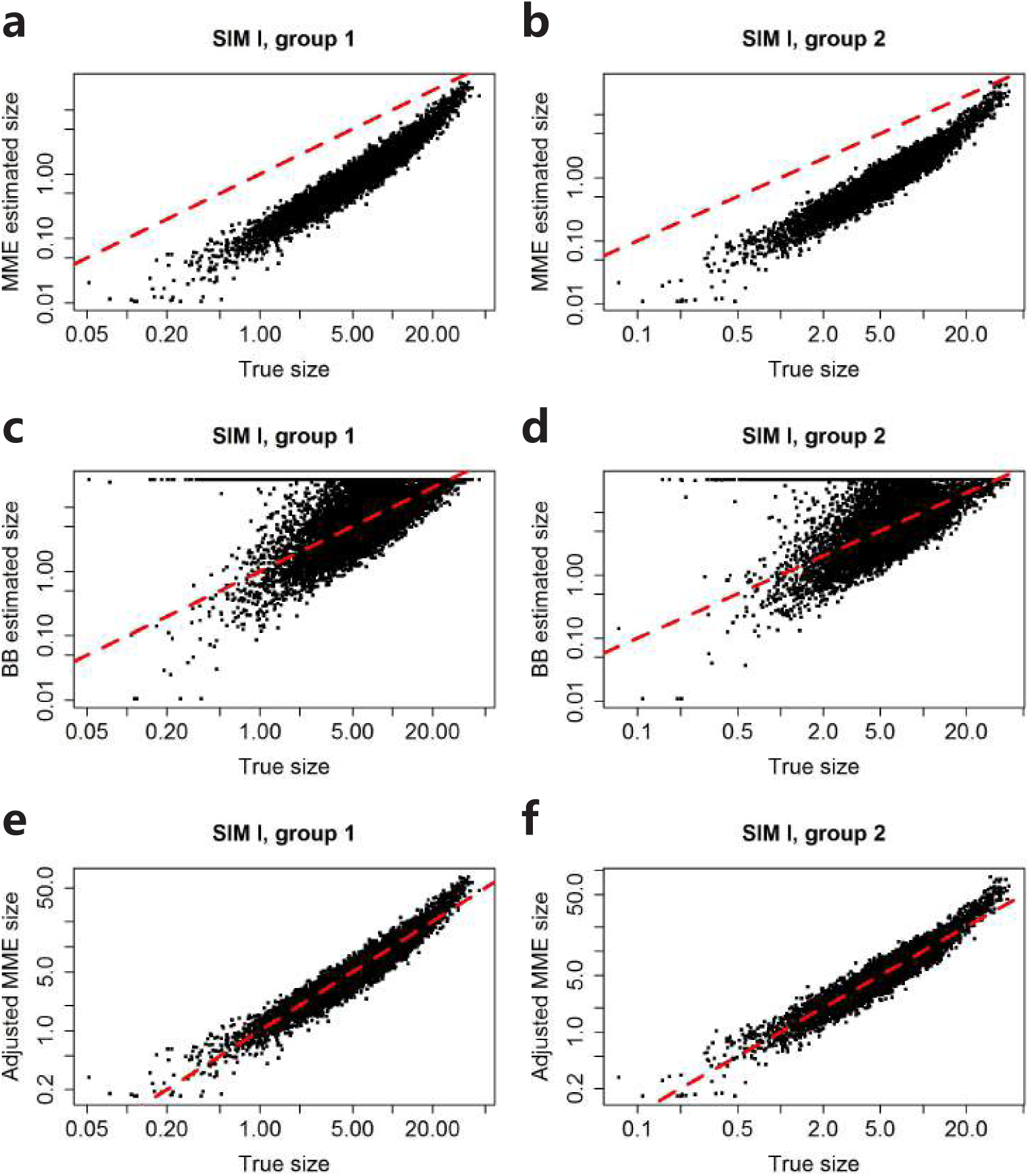
Estimation of the size factor (dispersion parameter) of the negative binomial prior distribution based on simulation studies (See Supplementary note 1 and 2). (a-b) comparison between the MME estimated size and the true size. (c-d) comparison between the BB estimated size and the true size. (e-f) comparison between the adjusted MME size and the true size. 2000 out of 10000 genes were simulated to be differentially expressed in group 1. Results are similar for other three simulated datasets (SIM DE II-IV).

## Supplementary Information

### Supplementary Note 1: two simulation protocols with Binomial distribution

#### “Binomial Splatter” simulation protocol

We adapted the simulation protocol proposed in the R package Splatter[4] but made two main modifications to that protocol:

1. We do not multiply the mean of the Gamma distribution by the library size factors. Instead, we add cell specific factors (capture efficiencies *β*_*j*_) at the last stage of simulation: Binomial step.

2. Unlike Splatter, we do not model the dropout rates explicitly. Instead we dropouts are the result of Binomial downsampling at the last stage of the simulation, which leads to a dropout vs mean expression relationship in the simulated data very similar to the one of experimental data (Supplementary Figures S2c, S3c, S4c, S5c and S6c).

The details of the simulation procedure are as follow:

1. We simulate a vector of base mean expressions 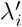 such that

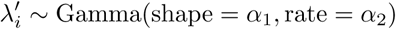

2. We simulate a vector of outlier factors *Ψ*_*i*_ such that *Ψ ∼* ln*𝒩* (*µ*^0^, *σ*^0^). For a proportion *π*^0^ of genes, we multiply the base mean expression by outlier factors: 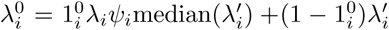, where 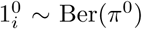. The above two steps are the same as those implemented in Splatter.

3. In the simulations with differential expression (SIM DE), the mean expression 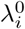 for the two groups are the same except that in the first group, we multiply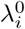 by a vector of DE factors simulated from the log normal distribution. Conversely, in SIM noDE study, no DE genes were simulated.

4. Then,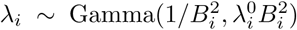, where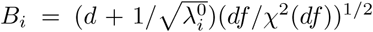 stands for the Biological Coefficient of Variation (BCV), and *d* is common dispersion. 1*/B*^2^ corresponds to the dispersion parameter *φ* in the Negative Binomial distribution.

5. Then the true count 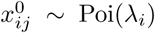. So far, no cell-specific factors have been taken into consideration.

6. Then a vector of cell specific capture efficiencies *β*_*j*_ needs to be specified in the simulation. When we compare the simulated data with the real data, the capture efficiencies estimated from the real data are used in the simulation. In all simulation studies, *β* is simulated from the log normal distribution and normalized to a specific mean capture efficiency (either 0.05 or 0.1).

7. Lastly, we implement the binomial step to obtain the observed count (binomial downsampling):

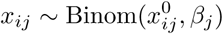

#### “Binomial bayNorm” simulation protocol

Unlike previous simulation protocol where gene expression parameters were simulated from a specific distribution with several estimated parameters, here we use gene specific priors estimated by bayNorm together with *β*_*j*_ to conduct gene and cell specific simulation. So, this method produces exactly simulated data of the same size as the real data (*P × Q*).

Let *µ*_*i*_ and *φ*_*i*_ be the estimated mean expression and dispersion parameter obtained by bayNorm for the *i*^th^ gene. Firstly a mean expression matrix 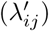 which is of the same dimension as the real data is created, such that 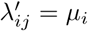across j. Then sampling from the following distributions leads to the simulated data:

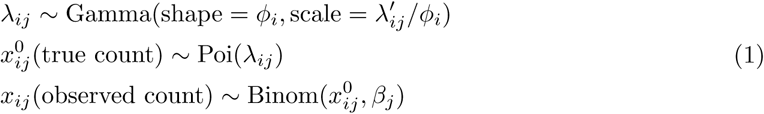

#### Parameter estimation from the real data (“Binomial Splatter” simulation protocol)

The parameter estimation methods used in the simulation are basically the same as those in Splatter, except that the input raw data are scaled by *β*_*j*_, before fitting the mean expression of the scaled data using Gamma distribution to estimate *α*_1_ and *α*_2_.

The estimation of other parameters like *π*^0^, *µ*^0^, *σ*^0^, *B*_*i*_ were achieved based on library size normalized data as also implemented in Splatter. Non-UMI based data were scaled and rounded before parameters were estimated as explained before.

### Supplementary Note 2: Simulation studies using the “Binomial Splatter” simulation protocol

We estimated parameters from the Klein and Bacher studies (92 H1-4M hESCs) and then generated 6 simulated datasets for comparing different normalization methods for their performance in correcting different capture efficiencies (study without DE genes), and in DE genes detection.

Two simulated datasets are generated without DE genes (homogeneous cells across two groups, but different mean capture efficiencies):

#### SIM noDE study I

(Using parameters estimated from the Klein study): The mean capture efficiencies of the two groups are 0.1 and 0.05 respectively. Simulated data was used in: Supplementary Figures S14, S18a-b, S19b.

#### SIM noDE study II

(Using parameters estimated from 92 H1-4M hESCs of the Bacher study): The mean capture efficiencies of the two groups are 0.1 and 0.05 respectively. Simulated data was used in: Supplementary Figures S15, S19c.

Simulation SIM DE studies are all based on parameters estimated from the Klein study.

#### SIM DE study I

The mean capture efficiencies for the two groups are both 0.1. Simulated data was used in: Supplementary Figures S12a,e,i, S17a, S18c,d, S21a and S24.

#### SIM DE study II

The mean capture efficiencies for the two groups are 0.05 and 0.1 respectively. Simulated data was used in: Supplementary Figures S12b,f,j, S17b, S18c,d, and S21b.

#### SIM DE study III

The mean capture efficiencies for the two groups are 0.1 and 0.05 respectively. Simulated data was used in: Supplementary Figures S12c,g,k, S17c, S18c,d, and S21c.

#### SIM DE study IV

The mean capture efficiencies for the two groups are both 0.05. Simulated data was used in: Supplementary Figures S12d,h,l, S17d, S18c,d, and S21d.

Genes with 0 counts across two groups were filtered out at the very beginning. No cells were filtered out. Belows are details about parameter settings used in the simulation studies which were estimated from the raw data as discussed above.

#### Parameters estimated from the Klein study

Most parameters were estimated from the Klein study except *β*. For each group, 10000 genes and 100 cells were simulated. The base mean expression for both groups were simulated from the Gamma distribution with *α*_1_ = 1.889 and *α*_2_ = 0.1229. *π*^0^ = 3% across two groups. The outlier factors were simulated from the log normal distribution with *µ*^0^ and *σ*^0^ set to 2.3 and 0.75 respectively. BCV was calculated with *d* = 0.12 and *df* = 105. The estimation of *β* is discussed in the Supplementary Note 3.

In the SIM noDE study I and the SIM DE studies I-IV, two groups of 100 cells with 10000 genes were simulated using the above parameter settings. *β*_*j*_ are simulated from the log normal distribution within each group with mean and sd (log scale) set to 2.74 and 0.3908 respectively. Within each group, we normalized the *β* to either 0.1 or 0.05.

In the SIM DE I-IV studies, the DE factors in the first group are simulated from the log normal distribution with log scale mean and sd set to 1 and 0.5 respectively.

#### Parameters estimated from the Bacher study (based on 92 H1-4M hESCs, 4 million mapped reads per cell)

Parameters were estimated from the raw data scaled by 20. *α*_1_ = 0.4129 and *α*_2_ = 0.005766. Outliers genes were simulated with *π*^0^ = 0.7%, *µ*^0^ = 4.745 and *σ*^0^ = 0.6027. BCV was calculated with *d* = 0.3113 and *df* = 7.6859.

In the SIM noDE case study II, two groups of 100 cells with 10000 genes were simulated using the above parameter setting. *β*_*j*_ are simulated from the log normal distribution within each group with mean and sd (log scale) to be -2.276 and 0.6886 respectively. Within each group, we normalized the *β* to either 0.1 or 0.05.

### Supplementary Note 3: Publicly available datasets and their preprocessing

#### Bacher study (non-UMI)

Single-cell RNA-seq expression data were downloaded from GEO GSE85917[5]. In this experiment, two groups of undifferentiated H1 hESCs were sequenced to a depth of 4 million mapped reads per cell and 1 million mapped reads per cell respectively. A similar experiment was done for H9 hESCs cells. The following filtering protocol was used in our study: spikeins and genes which do not have at least 10 non-zero counts were removed. After filtering, each group of H1 cells had 92 cells and 13181 genes, while groups of H9 cells had 91 cells and 13195 genes. Since these are non-UMI based data, we divided the raw data by a factor 20 for the 4 million mapped reads group and 10 for the other group so that scaled and rounded raw data were closer to the theoretical dropout vs mean curve (Supplementary Figure S9 a-d).

In order to estimate *β*, we let the total counts of observed spikeins in each cell be the scaling factors *s*_*j*_, and then normalized to 0.1 (based on scaled ERCC data, see Methods). As discussed in the text, within a 2 fold window of mean *β*, the performance of bayNorm in terms of DE detection is consistent (Supplementary Figure S20 a). Since cells in the two groups are the same, prior parameters were estimated across two groups when applying bayNorm (“global priors”).

Data was used in the Supplementary figures: S7, S9a-d, S13, S19a and S20a.

#### Islam study (non-UMI)

Raw data (48 ES cells and 44 MEF cells with 7284 genes) and a list of benchmark DE genes were kindly provided by Maria K. Jaakkola[6]. Genes which have zero expressions across all the 92 cells were removed in advance which left us with 5826 genes. Raw data was divided by a factor 10 before applying bayNorm on it as this is a non-UMI data and the scaled and rounded raw data is closer to the theoretical dropout vs mean curve.

For estimating *β*, scaling factors were estimated using scran[2] and were normalized to 0.03 (see Methods). The impact of different 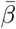 can be found in Supplementary Figure S20 b.

Data was used in the Figure 3c, Supplementary figures: S9e-f, S10e and S20b.

#### Patel study (non-UMI)

Raw data were stored in the Rpackage “patel2014gliohuman”https://github.com/willtownes/patel2014gliohuman[7]. Single cell data were scaled by a factor 20 and rounded as this is a non-UMI data and the scaled and rounded raw data is closer to the theoretical dropout vs mean curve. Cells with total counts less than or higher than the tenth percentile of total counts across cells were filtered out. In addition, genes with mean expression less than 1 were also removed. *β* were estimated using scran[2] with parameter “positive=TRUE”. The size factors were normalized to 0.06 (see Methods). Finally, cells with *β <* 0.01 were filtered out, leaving 590 cells and 5519 genes in the final datasets. We applied bayNorm on the scaled, rounded and preprocessed dataset. The estimated priors were used as input for Binomial bayNorm simulation protocol (Supplementary Figure S10f).

Data was used in the Supplementary figures: S10f.

#### Tung study (UMI)

Filtered molecule count matrix as well as the code for estimating *β* using spikeins were downloaded from the GitHub repository: https://github.com/jdblischak/singleCellSeq [8]. The list of benchmark DE genes was kindly provided by the author of R package DECENT[3]. Genes with 0 values across all three individuals were filtered out leaving 13058 genes in the final dataset. No cells were filtered out, resulting in 142, 201 and 221 cells for individuals NA19098, NA19101 and NA19239 respectively.

Data was used in the Figure 4, Supplementary figures: S3-5, S8c-d, S10b-d and S22-23.

#### Grün study (UMI)

Single-cell RNA-seq expression data were downloaded from GEO GSE54695[9]. The smFISH data used in that paper was kindly provided by the author.

The downloaded data was transformed to transcript number. We adapted the code provided in the supplementary material of [10] to convert the data to UMI count. We followed the same criterion as [10] for filtering genes in the 2i and serum data respectively. After filtering, we kept 74 cells and 9377 genes for the 2i medium data. For the serum medium data, we kept 52 cells and 9440 genes.

smFISH data were normalized by scaling factors which were calculated as cell sizes divided by mean of cell sizes.

Total number of input spikeins was estimated by adapting the code provided in the supplementary information of [10]. We divided the total number of observed spike-ins in each cell by the total number of input spike-ins to obtain scaling factors. We used smFISH data to estimate 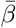for single cells under 2i and serum medium respectively (0.1212 and 0.1187 respectively, Supplementary Figure S11a-b). To obtain *β*, we normalized scaling factors (see Methods) to the corresponding 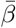 within each one of the two datasets.

Data was used in the Figure 2a,c,e,g and Supplementary figures: S11a-b.

#### Klein study (UMI)

Single-cell RNA-seq expression data were downloaded from GEO GSE65525[11]. ES cells data at day 0 (933 cells with 24175 genes) were used for simulations. We did not filter out any genes or cells.

For estimating *β*, trimmed mean of each cell at 1% was used and was normalized to 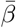set to 0.06[11] (see Methods).

Data was used in the Figure 1b-e, Figure 3a-b, Supplementary figures: S2 and S8a-b.

#### Torre study (UMI)

Single-cell RNA-seq expression data were downloaded from GEO GSE99330 (GSE99330 _dropseqUPM.txt.gz)[12]. This data matrix was converted to counts using a code kindly provided by the author of SAVER[13].

There are 32287 genes and 8640 cells in the raw data. First cells with less than 2000 genes detected or where the gene “GAPDH” could not be detected were filtered out. Second, genes with mean expression less or equal to 0.01 were removed. Third, the gene “GAPDH” was removed because it is used as a proxy for cell size[13] by normalizing it to the mean *β* which was estimated using smFISH data (see Methods). The final filtered dataset contained 9289 genes and 1134 cells.

For smFISH data, we filtered out cells with “GAPDH” counts below the bottom 10^th^ percentile or above the top 10^th^ percentile of ‘GAPDH” counts. smFISH data were then normalized by the expression of GAPDH divided by GAPDH mean expression[13].

We fitted a linear regression of the mean expression of filtered dataset on that of the normalized smFISH datset (Supplementary Figure S11c). The coefficient of explanatory variable was then used as 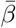 “GAPDH” expression, which was filtered out previously, was divided by the median and multiplied by 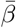 (see Methods).

Data was used in the Figure 2b,d,f,h, Figure 3a-b, Supplementary figures: S6, S8e-f and S10a.

#### Soumillon study (UMI)

The dataset was downloaded from GEO GSE53638 (GSE53638_ D3_ UMI.dat.gz for the single cell data and GSE53638_D3_Bulk UMI.dat.gz for the bulk data)[14]. DE detection was performed between the stage-3 differentiated cells at day 0 (D3T0) and day 7 (D3T7) (23895 genes and 1949 cells) [3]. Cells with library sizes below the bottom and above the top 5^th^ percentiles were filtered out. Genes with mean expression across two groups greater than 0.05 were retained, resulting in a dataset of 1754 cells with 8586 genes (832 cells and 922 cells belonging to day 0 and day 7 time-points respectively). Using the same 8586 genes in the bulk dataset, the reference DE genes were defined to be the top 1000 genes which have the greatest log fold-change in the corresponding bulk RNA-seq data[3].

Based on the smFISH data, 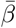 is expected to be in the range of 1_2%[14]. Here we set 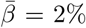. Scaling factors were estimated using R package scran[2], and then normalized to 0.02 (see Methods).

Data was used in the Figure 3d and Supplementary figures: S16.

### Supplementary Note 4: Normalization methods and relevant R packages

#### Splatter, R package version 1.4.1

For both UMI and non-UMI data, the default settings of Splatter were used for estimating parameters from the input data and simulating scRNAseq data.

For non-UMI data in Supplementary Figure S7, H1 P24 single cell data were divided by 20 and then rounded as an input for Splatter.

#### SAVER, R package version 0.4.0

We used the default settings for SAVER throughout the paper. When estimating mean, CV and Gini coeffieicnts, for both bayNorm and SAVER we generated 5 samples for the Torre study and 20 samples the Gruün study (3D arrays). The mean, CV and Gini were estimated across the cells and samples. In the 6 simulation studies, 10 samples were generated from posterior for both bayNorm and SAVER. In the Klein (Figure 3a-b), Tung and Soumillon studies, 5 samples were generated. In Tung study, SAVER was applied within each individual.

#### SCnorm, R package version 1.1.0

We used the default settings in SCnorm except for UMI datasets and simulated data, where “dither Counts=TRUE” was used as UMI data contains tied counts.

#### scImpute, R package version 0.0.6

We applied scImpute using its default settings. In the Tung study, scImpute was applied on each individual independently.

#### MAGIC, R package version 0.1.0

MAGIC was applied using its default settings. In the Tung study, MAGIC was applied within each individual.

#### DCA, Python package version 0.2.2

DCA was applied using its default settings. In Tung study, since genes with 0 counts across cells within each individual could be filtered out, we applied DCA acorss all cells.

#### Scaling method

Throughout the paper, the scaling method refers to the modified formula **??** such that 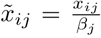 In UMI datasets and simulated data, *β* used in scaling method are as the same as that used in bayNorm.

For the Bacher study (non-UMI), the scaling factors were set to be the total counts of spike-ins normalized to 0.1 (see Methods).

For the Islam study (non-UMI), the scaling factors were estimated based on raw data using R package scran with “sizes=c(20,30,40,50)” and “positive=TRUE”, which were further normalized to (see Methods).

